# Extracellular matrix properties and dynamics are important for butterfly scale nanostructure development

**DOI:** 10.64898/2026.09.22.753460

**Authors:** Anupama Prakash, Victoria Lloyd, Yacine Ben Chehida, Veronika Shkaraburova, Rachel George, Trong Khoa Pham, Mark O. Collins, Sam Amsbury, Nicola J. Nadeau

**Affiliations:** School of Biosciences, University of Sheffield, Sheffield, United Kingdom; biOMICS Mass Spectrometry Facility, University of Sheffield, Sheffield, United Kingdom; Natural History Museum, London, United Kingdom; Leverhulme Centre for Anthropocene Biodiversity, Department of Biology, University of York; York, United Kingdom

## Abstract

Butterfly wing scales are intricately structured, cuticle-derived, extensions of single scale cells. Their highly ordered structures confer a range of functions, including producing structural color, which depends upon unknown molecular mechanisms controlling the secretion and ordering of an extracellular matrix (ECM). To find the genes required for patterning the ECM we compare scale nanostructure development between structurally colored blue scales and non-iridescent black scales on the forewings of the butterfly *Heliconius sara* using transmission electron microscopy, RNA-seq and proteomics. We first determined that structural differences between the two scale types are established from about half-way through pupal development and then investigated differences in relevant genes and proteins between scale types from this stage onwards. We show that knockouts of the genes coding for zona pellucida (ZP) domain proteins, *miniature* and *trinity*, and the cuticle protein gene, *Cpr128* produce distinct effects on scale morphology and structure formation. In particular, some highly unusual scale morphologies are observed in *miniature* knockout individuals, suggesting a previously undescribed role of ZP domain proteins in structuring the ECM of scale cells and in mediating the mechanochemical interactions needed to produce the complex extracellular structures important for biological function in this system.

## Introduction

Many cells produce hierarchically structured extracellular materials to serve multiple functions from anti-reflectivity, hydrophobicity, molecular filtering, structural support, sound production, or color (Yadav and Majumder 2021; Gorb 2009). They vary from insect cuticles over the body and wings (Noh et al. 2016; Moussian 2010; Tajiri 2017; Duan et al. 2026), tracheal folds (Öztürk-Çolak et al. 2016), hairs on human skin, ridges on flower petals and leaves (Surapaneni et al. 2020; Wilts et al. 2018), bristles, and scales (Ghiradella 2010) to name a few. In each case, the cells secrete a complex extracellular matrix that is shaped by an intricate coupling of biomechanical forces on the cell/tissue and cytoskeletal dynamics within each cell. Extracellular matrices in the context of insect cuticles are not single homogenous materials, but multi-layered combinations of various biomolecules secreted and assembled sequentially (Ghiradella 1974; Moussian 2010; Moussian et al. 2006; Noh et al. 2016; Duan et al. 2026).

Lepidopteran wing scales are complex chitinous structures produced by single cells (Dinwiddie et al. 2014; Ghiradella and Butler 2009). The scale precursor cell divides into two daughter cells; one that becomes the future scale and the other that produces a socket, attaching the scale to the wing membrane (Dinwiddie et al. 2014; Prakash et al. 2024; Loh et al. 2025). Extension of the apical cell membrane of the scale cell through the socket, creates an initial cytoplasm filled scale bud. This extension further grows and flattens to create two laminae. Secretion, organization and hardening of various extracellular matrix components outside the cell membrane produces the future adult scale (Ghiradella 1994; Locke 1966). In a typical scale structure, the lower lamina is most often a thin-film while the upper lamina is intricately sculpted into ridges and cross-ribs, connected to the lower lamina via pillar-like trabeculae in the scale lumen. Wing scales can also have pigments in the chitinous matrix that create pigmentary colors and/or have nanostructures in different scale sub-structures that produce structural colors by the constructive interference of certain light wavelengths (Giraldo and Stavenga 2016; Prakash et al. 2022; Stavenga et al. 2014). Scale nanostructures can vary from intra-lumen structures like photonic crystals to ridge nanostructures based on overlapping ridge lamellae (Thayer and Patel 2023).

Despite a good understanding of the various biochemical processes and genes in pigment pathways of scale cells (Matsuoka and Monteiro 2018; Hines et al. 2012), knowledge about the genetics, development and biomechanics of scale nanostructures remains poor. One of the important players, the cytoskeletal protein actin, determines scale shape at multiple developmental stages (Dinwiddie et al. 2014; Lloyd et al. 2024; Seah and Saranathan 2023). Initial constraints provided by actin bundles within the newly developed scale bud determine ridge positions and lead to plasma membrane buckling that form proto-ridges (Totz et al. 2024). Further, actin re-organization during scale development associates with the formation of various scale sub-structures (Lloyd et al. 2024; Seah and Saranathan 2023). However, how ridges grow and elaborate to form lamellae and micro-ribs, how other sub-structures like cross-ribs and trabeculae arise, as well as the functions of numerous extracellular matrix components like proteins, lipids and chitin in scale structure determination is still largely unknown (Prakash et al. 2026).

To better understand how extracellular cuticle secretion dynamics contribute to scale structure formation, we investigate the development of scale nanostructures in the butterfly *Heliconius sara*. We compare iridescent structurally colored blue scales to non-iridescent black/brown scales, both scale types being highly pigmented (Parnell et al. 2018; Wilts et al. 2017). We first characterize the temporal changes in extracellular matrix formation between the two scale types. We then perform comparative RNA-seq and proteomic analyses over a time series of pupation and test the spatial expression of various candidate genes using fluorescent *in situ* hybridization and immunohistochemistry, before functionally testing the role of candidate extracellular matrix genes in determining scale shape using CRISPR/Cas9 targeted gene editing. Together, these findings emphasize the role of extracellular matrix properties and dynamics in structural color development.

## Results

### Differences in epicuticle folding establish early nanostructural differences in wing scales

*H. sara* butterflies exhibit clearly defined dorsal forewing regions with either blue, yellow, or black wing patches. Both blue and black scales are highly pigmented and appear black at certain angles of observation. However, scales at the basal region of the forewing are iridescent blue (Fig 1A (i)). To explore the differences in scale nanostructure that produce iridescent vs non-iridescent colors we imaged adult scales using both scanning and transmission electron microscopy (Fig 1A, B). Adult blue scales comprised closely spaced longitudinal ridges (Lloyd et al. 2024) that were relatively smooth throughout their length. From an angled view, these scales had two parallel lamellae running along the ridge with smaller microribs below (Fig 1A (i), red arrows). In comparison, adult black scales had ridges spaced farther apart (Lloyd et al. 2024)with the ridges broken into overlapping ridge lamellae (Fig 1A (ii), red arrows). Parallel lamellae were absent in these scales and microribs extended from the top of the ridge.

**Figure 1:**
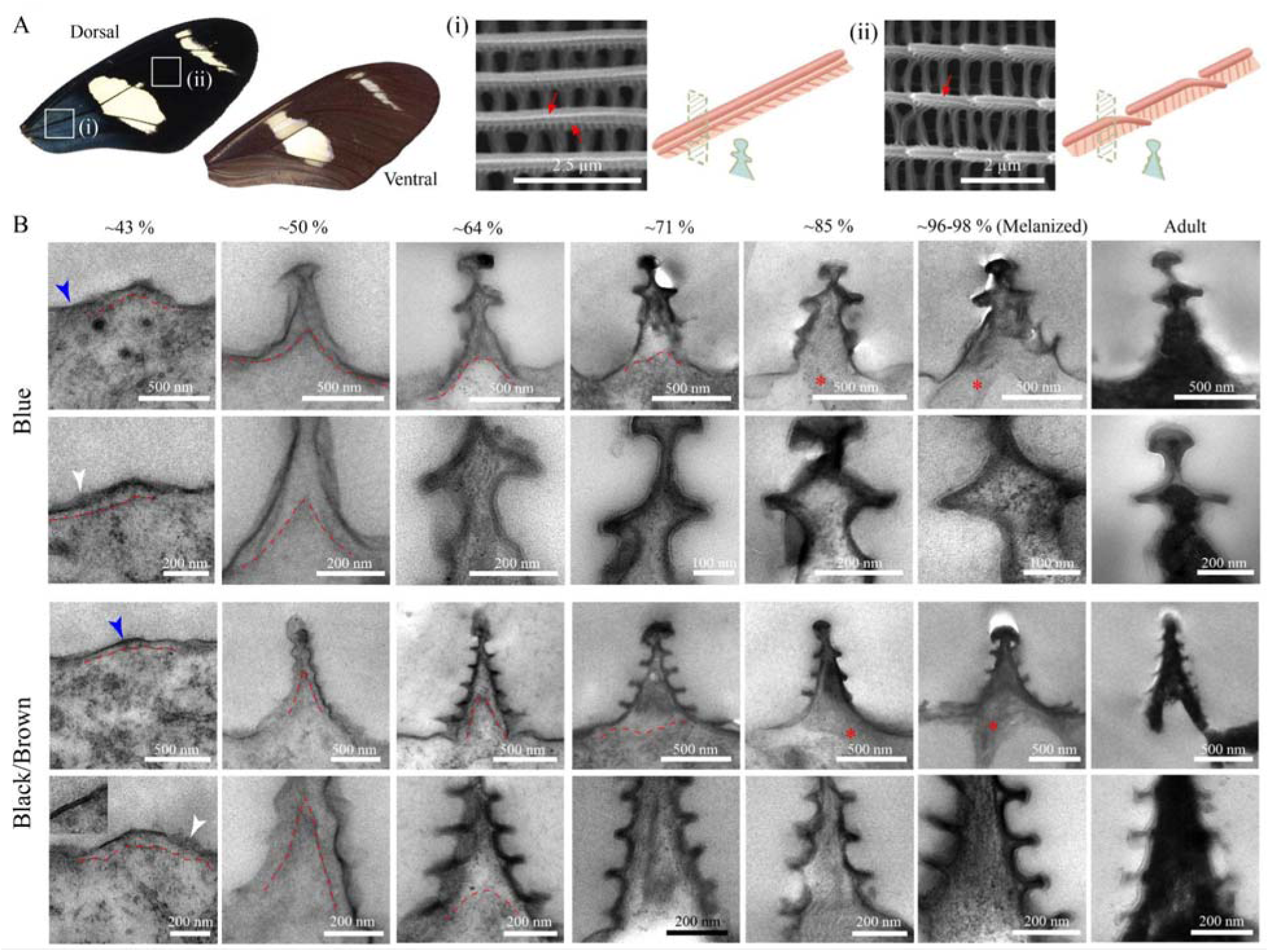
Differences in epicuticle folding establish early nanostructural differences in *Heliconius sara* wing scales. (A) *H. sara* dorsal forewing patterned by three blocks of color. (i) and (ii) are scanning electron microscopy images of iridescent blue scales and pigmented black scales respectively. Red arrows mark the smooth parallel lamellae in the blue scale ridges and the broken overlapping ridge lamellae in the black scales. The schematics illustrate the structure of a single ridge in each scale type and its cross-section. (B) A time-series of transmission electron microscopy cross-sections of a ridge in an iridescent blue scale (top) and black/brown scales (bottom). For each set, an entire ridge cross-section is shown at the top and a magnified image is presented below. Dashed red lines mark the plasma membrane of the scale cell and red stars indicate trabeculae. Blue arrowheads point to the epicuticular envelope, and the white arrowheads point to regions of the envelope that are still developing and appear fuzzy.

To understand the development of these different scale nanostructures and pinpoint the timing of their structural bifurcation, we performed a TEM time series over pupal development (PD), comparing scale cross-sections from the blue regions of the forewing to black/brown scale cross-sections. At the very early stages, ~28 % PD, scales were growing out from the wing membrane via the extension of the apical cell membrane of the scale cells (Supp Fig 1). By ~43 % PD, the initial cylindrical scales had flattened. Outside the cell membrane, there was a membrane-proximal zone, and the formation of an electron-dense envelope was visible at the edge of this zone (Fig 1B-43 %; blue arrowheads, Supp Fig 2). The double-layered membrane (Inset, Figure 1B-43 %) is the first layer of the extracellular cuticular matrix to be synthesized (Ghiradella 1974; Moussian et al. 2006). The formation of the envelope was not uniform across the cell and appeared to self-organize in patches as seen from fuzzy regions interspersed between organized double membrane regions (Fig 1B-43%; white arrowheads, Supp Fig 2). This was similar to envelope formation during cuticle development in other species (Moussian et al. 2006). No visible differences were present between blue and black scales up to this point.

At 50 % PD, blue and black scale cross-sections differed in their shape. Though both scale types had developed the bulge at the top of each ridge (the ridge lamella) by a buckling of the envelope, the envelope of blue scales appeared smooth while the envelope of the black/brown scales was wavy (Fig 1B). By 64 % PD, these differences were exacerbated with the blue scales developing the parallel lamella under the top of the ridge. This was visible as buckled extracellular matrix (ECM) on either side like two extended arms. The black/brown scales at this point lacked arm-like extensions and instead displayed numerous smaller buckling of the ECM. In both cases the ECM had thickened and included an electron dense layer ~10 nm thick, below the outer envelope. We call this set of layers the epicuticle, based on similarities with *Drosophila melanogaster* cuticle development (Moussian et al. 2006) and references to this layer in previous butterfly literature (Ghiradella 1974; Locke 1966). In addition, the space enclosed by the epicuticle was now filled with an amorphous material secreted by the scale cell, the developing procuticle.

Between 65-85 % PD, development of the procuticle continued with the likely addition of various procuticle components including the protein matrix and chitin fibers (Fig 1B). Apart from the amorphous material filling up the ridges, no other changes were visible in the TEM. Around 85 % PD, the scale cell is retracting (Supp Fig 3A) and pillar-like trabeculae that would connect the upper and lower laminas in the adult scales begin to form (Fig 1B-85 %, red stars). The trabeculae appeared to be growing from either of the laminas towards the other or bi-directionally (Supp Fig 3A), though it’s hard to say if this is an artefact of the sectioning angle or if trabeculae can grow in either direction.

Black pigmentation occurs towards the end of pupal development in *H. sara*, only a couple of hours before emergence. Cross-sections of melanized blue and black scales at ~96-98 % PD indicated that the cell had fully retracted by this time point (Fig 1B, Supp Fig 3B-D). Melanization did not appear to make TEM cross-sections of scales highly electron-dense and the procuticle still appeared lighter than the electron-dense epicuticle. In the hardened adult scales however, the bulk of the ridges appears electron-dense surrounded by an electron-lucent material (Fig 1B, Supp Fig 4).

### Cuticle and ECM proteins are differentially expressed between blue and black tissues at later stages of pupal development

To identify genes and proteins that may be controlling the morphological differences we observed between the blue and black wing regions, we performed comparative RNA-seq and mass spectrometry-based proteomic analysis of the two tissue types across different stages of pupal development (Fig 2A). In both data sets, the main axis of variation separated the developmental stages (explaining 39% and 42% of the variation in RNAseq and proteomics respectively, Supp Fig 5). Later developmental stages became more similar, demonstrating increased stabilization as the wing nears maturity. Nevertheless, in both data sets there was an axis of variation that separated the blue and black wing regions (Supp Fig 5; Principal Component 5, representing about 3% of total variation in both cases), demonstrating that there are some consistent differences in expressed genes/proteins between wing regions through development.

**Figure 2:**
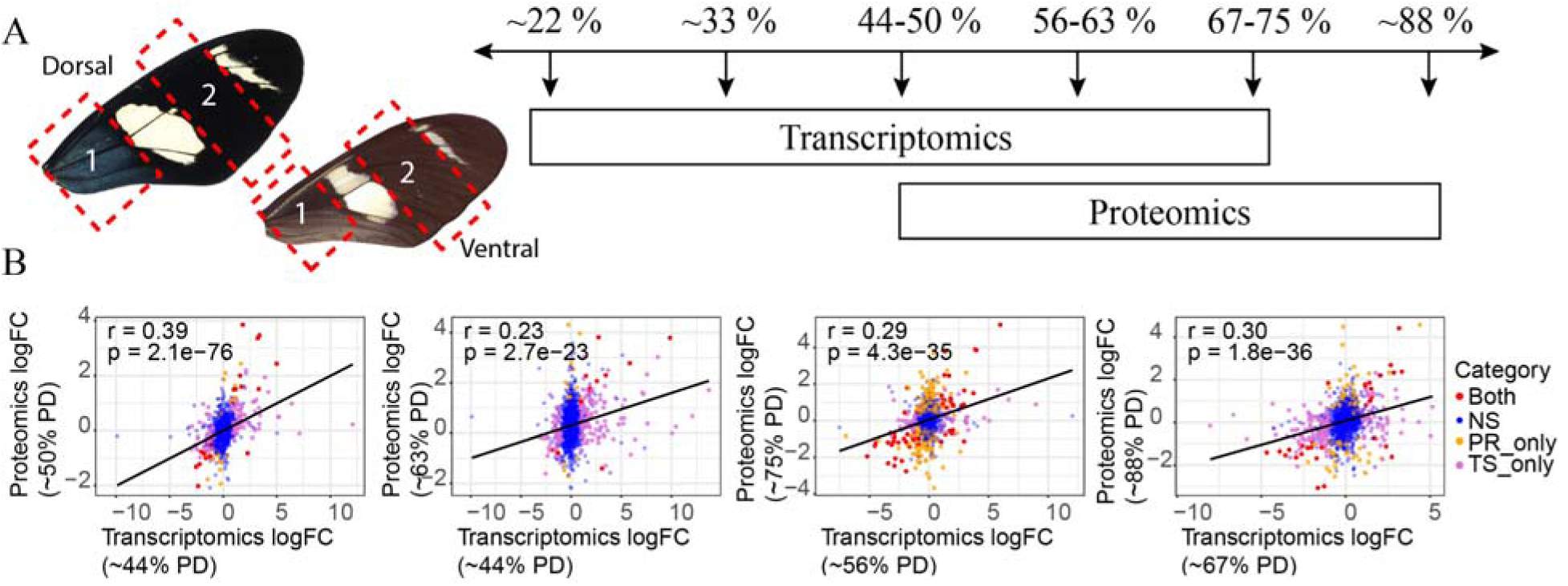
Differences in gene expression and protein abundance between blue and black wing regions are correlated at comparable developmental stages. (A) Blue (1) and black (2) tissue regions of the forewing dissected for both RNA-seq and proteomics analysis at various time points across pupal development (PD; measured as % PD). (B) Comparison of the relative expression (log fold-change) in blue versus black regions between data types at the stages showing the strongest correlations. Points are genes colored by whether they are DE (FDR<0.05) in both data sets (red), in the proteomics only (yellow), in the transcriptomics only (pink) or in neither (blue).

We found increasing numbers of differentially expressed (DE) genes and proteins between presumptive blue and black wing regions as development progressed (Fig 2, Supp Fig 6). Comparing the transcriptomic and proteomic data sets, at the earliest sampled time-point in both comparisons (4 days post pupation = 44-50% PD) there was a significant correlation between observed blue:black log-fold-change values (r = 0.39, p < 0.001, Fig 2, Supp Fig 7). At later developmental stages, the strongest correlations between the transcriptomic and proteomic comparisons became offset by a day, such that there was greater similarity in the DE genes between the earlier stage in the transcriptome with the later stage in the proteome (Fig 2, Supp Fig 7). This is consistent with there being a time-delay between the greatest difference in transcription and that difference appearing in the transcribed proteome.

Gene Ontology (GO) enrichment analysis of the differentially expressed genes and proteins showed significant enrichment in molecular functions and biological processes associated with the cuticle, ECM and cell adhesion (Supp Tables 1 and 2). DE of cuticle-related genes was particularly pronounced from ~44% PD and proteins from ~75% PD. There was also significant enrichment of DE genes associated with DNA binding and gene regulation in the earlier stages of the transcriptomic data, likely corresponding to transcription factors important in specifying the identity of the wing regions and scale types.

We identified genes that were differentially expressed and abundant in the transcriptomic and proteomic data sets respectively, and with predicted functions that made them interesting candidates for further investigation. Cuticular proteins are needed for correct scale structure formation in moth scales (Liu et al. 2021), making them promising candidates for nanostructure development in butterfly scales. We identified four cuticular proteins that showed strong and consistent DE in both proteomics and transcriptomics. CPR100a and CPR129 were both generally upregulated in blue compared to black wing regions, while CPR128 and LCP65Ac were both generally downregulated in blue compared black wing regions (Figure 3).

**Figure 3:**
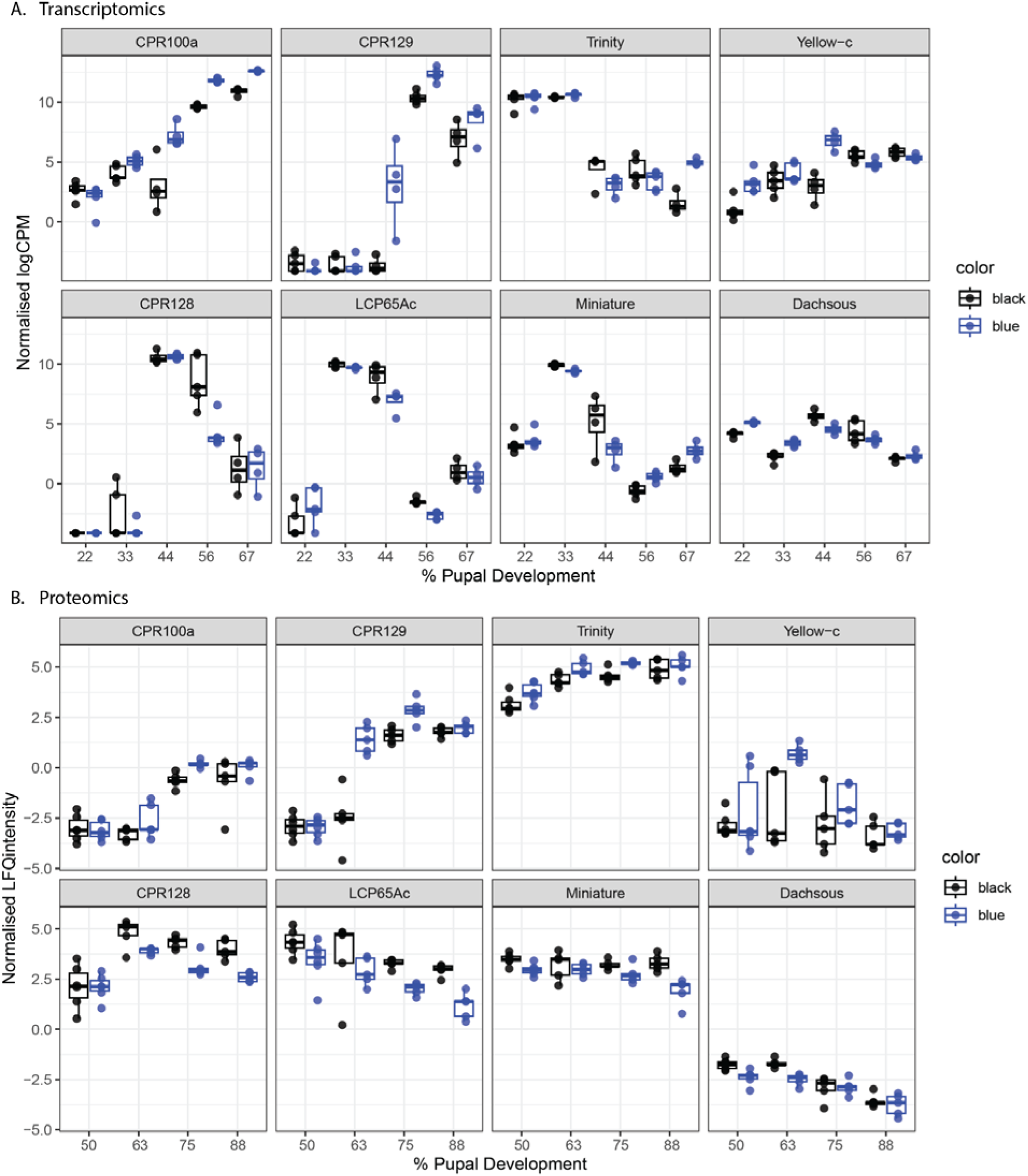
Eight candidate genes/proteins for control of scale structure show differential expression between blue and black wing regions. (Expression level as normalised counts per million (CPM) for the RNA-seq (A) and label-free quantification (LFQ) intensity for the proteomics (B) at different developmental stages (given as % pupal development) and for each tissue. These include four cuticle proteins (CPR100a, CPR129, CPR128 and LCP65Ac), two ZP domain proteins (Trinity and Miniature), Yellow-C and the cell adhesion protein Dachsous. Points represent values from individual samples and box-and-whiskers give the interquartile ranges, maxima and minima estimates for each sample group.

We also identified the gene/protein Yellow-c as being generally upregulated in blue compared to black wing regions (Figure 3). Yellow genes are involved in cuticle melanisation and maturation, and mutations in yellow have previously been found to alter scale structure (Matsuoka and Monteiro 2018). In addition, we identified the cell adhesion protein Dachsous as being generally up regulated in black compared to blue wing regions (Figure 3). This is an interesting candidate because the *Drosophila* orthologue is involved in controlling epithelial contraction and morphogenesis (Bosveld et al. 2012).

ZP matrix proteins were also particularly exciting because they are expressed in cuticle forming tissues and have previously been identified as important regulators of apical membrane shape, interacting between the ECM, membrane, and cytoskeleton (Roch 2000, Fernandes 2010). The ZP domain protein Trinity was consistently present at higher levels in the blue compared to black scales throughout development. Its expression was not quite as consistent in transcriptomic data but at the latest developmental stage (~67% of development) the transcript was significantly upregulated in the blue scales. Miniature is another ZP domain protein that differed significantly between wing regions, this time being more abundant in the black wing region. This difference was again less consistent in the transcriptomic data but at 38-50% PD, when its expression was also highest, it was more highly expressed in the black wing region (Figure 3).

### Cuticular protein CPR128 is expressed in scale/socket pairs and may function in cuticle formation

We first tested six candidates for their role in scale development: *yellow-c*, *Cpr129, Cpr100a*, *Lcp65A*, *Cpr128* and *dachsous*. The mRNA of two of the cuticle proteins, *Cpr129* and *Cpr100a* was strongly expressed in tracheal cells (Supp Fig 8) while *yellow-c* was highly expressed in the male androconial tissue, sensory cells at the base of the wing and the large, thick bristles that develop along the veins and margin (Supp Fig 9). No expression was seen in the scale cells so we did not proceed with functional tests for these genes.

*Lcp65A* mRNA was expressed in scale cells and had a marginally stronger expression in the black scale cells on the forewings (Supp Fig 10), consistent with the RNAseq and proteomics data. Dachsous expression, investigated using an anti-ds antibody, was strong in the hindwing androconial tissues (Supp Fig 11). Expression was also higher in the distal parts of both fore- and hindwings corresponding to the black wing regions (Supp Fig 11).

Interestingly, *Cpr128* mRNA appeared in neat rows corresponding to the scale cells at ~57% PD and was higher in the distal wing regions on both the dorsal and ventral surfaces, corresponding to black and brown scales respectively (Fig 4A), consistent with the RNAseq and proteomics data. Upon closer inspection, the mRNA at this time point appeared to be localized to the region just below the body of the external socket (Fig 4C, D), within the socket cell (Fig 4B; yellow arrowhead). However, CRISPR-Cas9 knockouts of *Cpr128* did not produce any noticeable defects in the sockets, which developed normally. Instead, some scale features, like the organization of the ridges and lamellae thickness were visibly affected in one crispant (Fig 4E, F, G), with effects also seen in the proximally located iridescent blue scales (Fig 4G).

**Figure 4:**
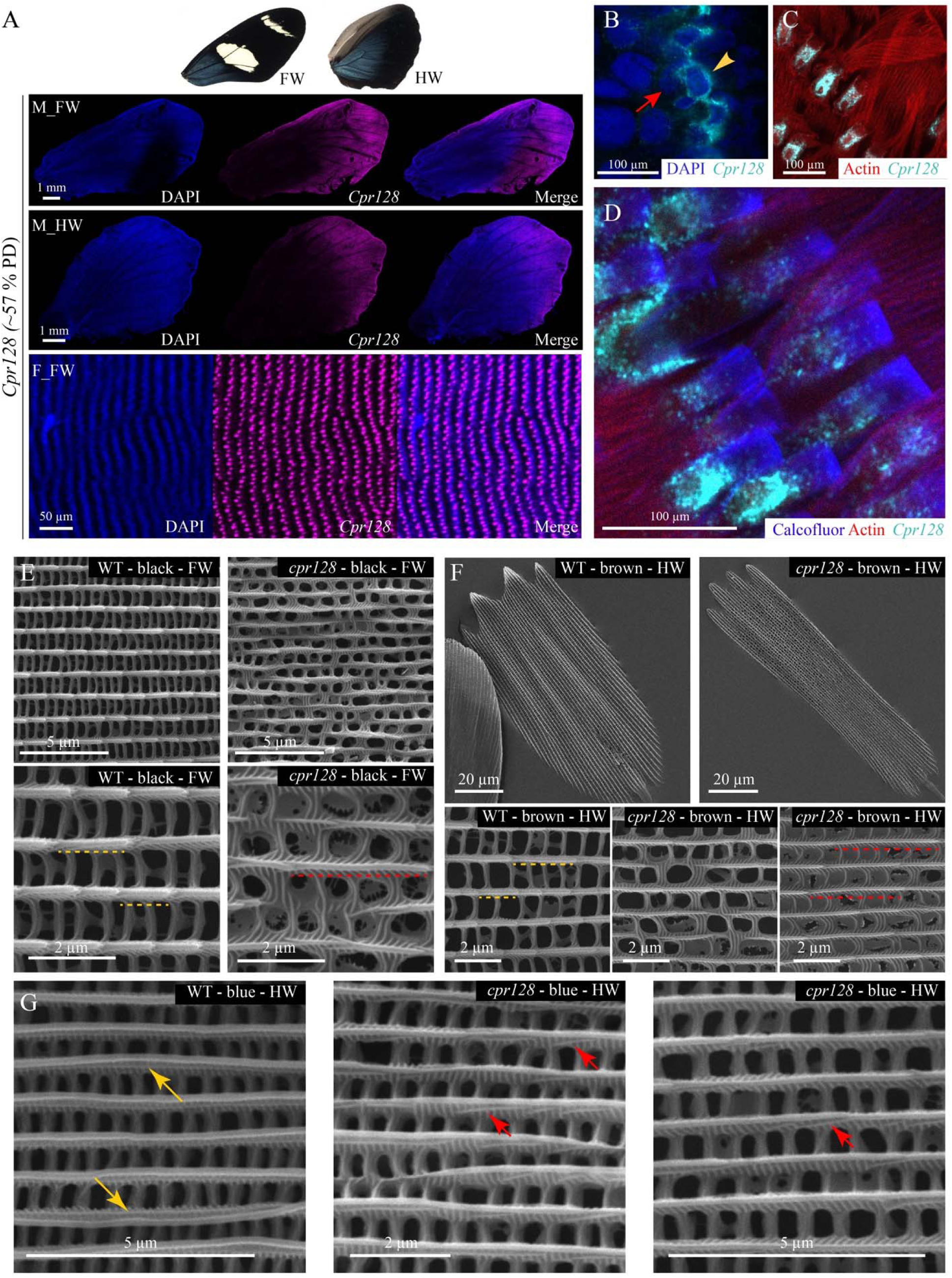
*Cpr128* mRNA is expressed in socket cells of the distal wing at 57% PD but demonstrates functional effects on scale formation. (A) At ~57% PD, *Cpr128* mRNA is expressed in neat rows corresponding to scale and socket cell pairs at the distal ends of both fore- and hindwings. (B) The nuclei of scale and socket cell pairs stained with DAPI (dark blue). *Cpr128* mRNA (cyan) is strongly expressed in the socket cell cytoplasm (yellow arrowhead) and is not visible in the scale cell (red arrow) (C, D) *Cpr128* mRNA visualized in the socket cells surrounding the extended scales projecting through them. Actin is marked in red; cuticle is blue and stained with calcofluor; *Cpr128* mRNA is in cyan. (E) Dorsal forewing black scales of a WT individual compared to those from a Cpr128 crispant. (F) Ventral hindwing brown scales of a WT individual compared to those from a Cpr128 crispant. In E and F, yellow dotted lines indicate lengths of some of the overlapping ridge lamellae in WT scales while red dotted lines indicate the much longer lengths of overlapping ridge lamellae in the Cpr128 crispant. (G) Dorsal hindwing blue scales of a WT individual compared to those from a Cpr128 crispant. Yellow arrows point to the unbroken, parallel lamella seen in WT blue scales. Red arrows point to the broken and often curved second lamella layer in the blue scales of the Cpr128 crispant.

### Extracellular matrix proteins *trynity* and *miniature* are crucial for correct cuticle formation in wing scales

We then investigated the two ZP domain proteins, *trynity (tyn)* and *miniature (min)*. Detailed investigation of *tyn* mRNA expression with fluorescent *in situ* HCR revealed that at ~30-40% PD it was expressed in scale cells but appeared to be uniformly expressed across the entire wing (Fig 5A), contrary to the RNA-seq results. Within the scale cells *tyn* mRNA was strongly polarized, appearing as a crescent shape, with the convex side towards the distal wing edge (Fig 5B). At ~50-57% PD, *tyn* mRNA was no longer expressed in the scale cells and was strongly localized to cells of the peripheral tissue, outside the future adult wing margin (Fig 5C).

**Figure 5:**
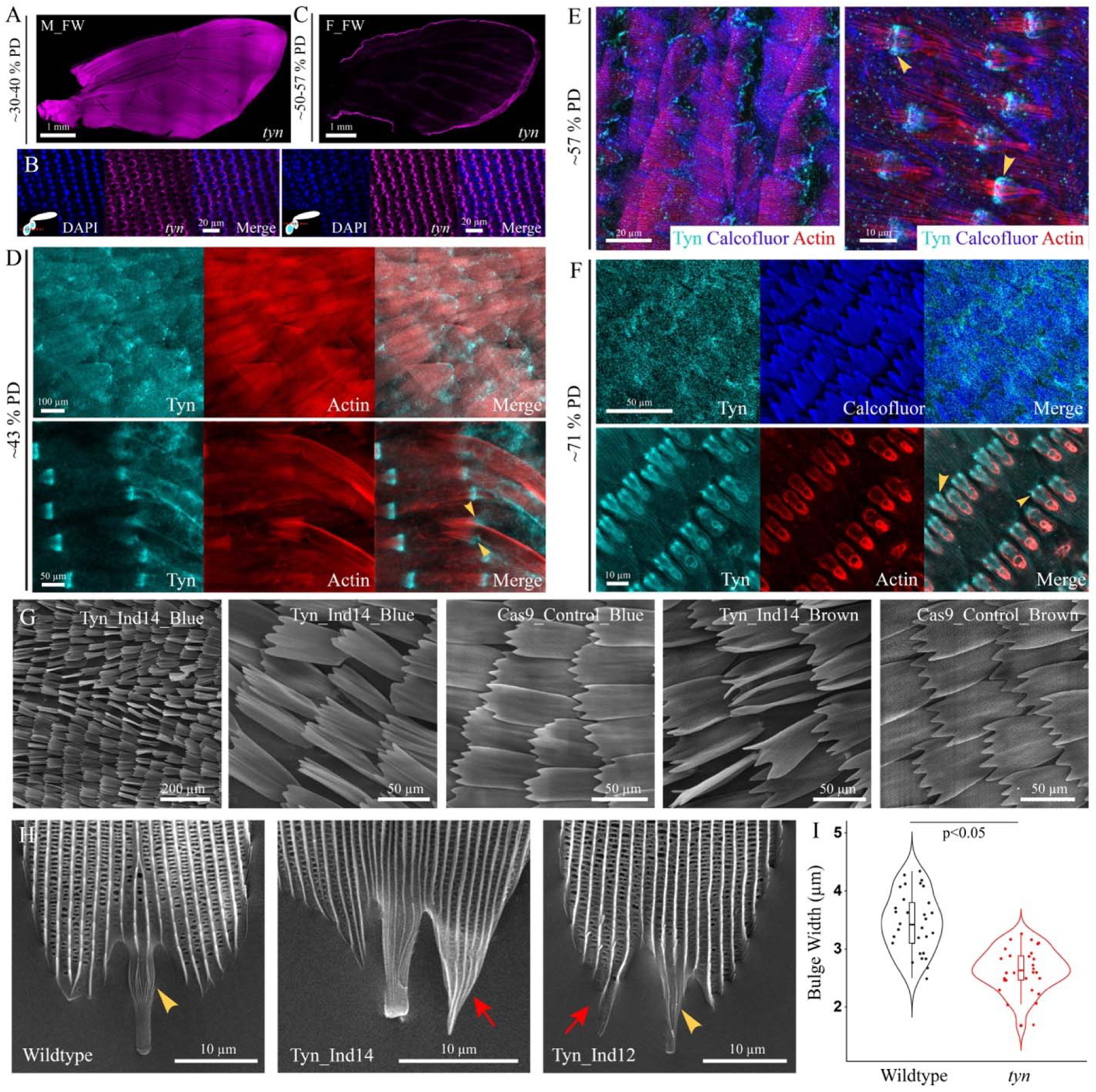
Trynity is needed for scale development and determination of basal scale features. (A) Between 30-40% PD, *tyn* mRNA is expressed in all scale cells. (B) *tyn* mRNA in scale cells is polarized with the convex side towards the distal wing edge. Two different z-planes as marked in the inset as shown. (Left) Towards the top of the wing, where the socket nuclei lie and (Right) deeper into the wing tissue where the larger scale nuclei lie. (C) Between 50-57% PD, *tyn* mRNA is no longer seen in the scale cells and is instead strongly expressed in cells of the peripheral tissue. In (A-C) *tyn* mRNA is in magenta and DAPI marks the nucleus in blue. (D-F) Tyn protein expression at (D) ~43% PD (E) ~57% PD and (F) ~71% PD in the scale blade and near the scale base. Tyn protein is in cyan, actin is marked in red and the plasma membrane is marked by calcofluor in blue. Yellow arrowheads indicate the strong Tyn extracellular protein expression at the scale base. (G) Scanning electron micrographs of *tyn* crispant and Cas9 control scales. (H) *tyn* crispant scale bases compared to a wildtype scale base. Yellow arrowhead marks the bulge in the scale stalk where it inserts into the socket, and the red arrows indicate the pointy “tail”. (I) Violin plot of the bulge width of wildtype and *tyn* crispant scales.

Using an antibody against Tyn, we were able to further investigate Tyn protein expression in developing scales and sockets. Between 40-70% PD, Tyn expression over the developing scale blade was not clear cut, initially appearing as a hazy layer over the scales (Fig 5D) and then becoming more punctate (Fig 5E, F, Supp Fig 12). However, Tyn expression at the base of the scales was well defined, appearing as a clear, extracellular band where the scales exited the sockets (Fig 5D, E; yellow arrowheads, Supp Fig 12), as seen in *Drosophila* mechanosensory bristles (Itakura 2026). Over time, Tyn proteins were concentrated extracellularly at this junction leading to scale constriction at the base and the formation of a small bulge of the scale stalk within the socket (Fig 5F, yellow arrowheads, Supp Fig 12).

CRISPR-Cas9 knockouts of *tyn* affected scale development across the wing, irrespective of color, leading to curved and malformed scales (Fig 5G). In many of the twisted scales, mosaic patches of unformed or improperly formed upper lamina nanostructures were visible (Supp Fig 13). These mosaic patches displayed incorrectly structured ridges and covered inter-ridge spaces without defined crossribs (Supp Fig 13). The scale stalk was affected, with loss of the bulge in many crispant scales (Fig 5H, yellow arrowheads). Scale bulge width differed significantly between wildtype and *tyn* crispants (F_1,59_ = 53.57, p < 0.0001). Wildtype scales had an average bulge width of 3.46 ± 0.08 μm (mean ± SE). *tyn* crispant scales had significantly narrower bulges, averaging 2.64 ± 0.08 (a reduction of 0.82 ± 0.11 µm, t_59_ = −7.32, p < 0.0001) (Fig 5I). Some scales also had pointy scale “tails” instead of the normally curved and rounded bases (Fig 5H, red arrows), though log-transformed scale tail length did not differ significantly between wildtype and *tyn* crispants (F_1, 52_ = 0.31, p = 0.579).

Similar to *tyn*, *min* also appears to have multiple functional roles across different cell types based on its temporal expression patterns (Fig 6A, B). *min* mRNA was expressed across the entire wing blade in the smaller epithelial cells at ~30% PD (Fig 6A). At a later stage of pupal development, *min* mRNA was no longer highly expressed in the epithelial cells but instead in the scale cells (Fig 6B) and more strongly in the black wing regions compared to the iridescent blue at ~43% PD (Fig 6B). Between ~43-70% PD, Min proteins were present in the scale blades as puncta, concentrating at the finger-like scale tips (Fig 6C-F, Supp Fig 14).

**Figure 6:**
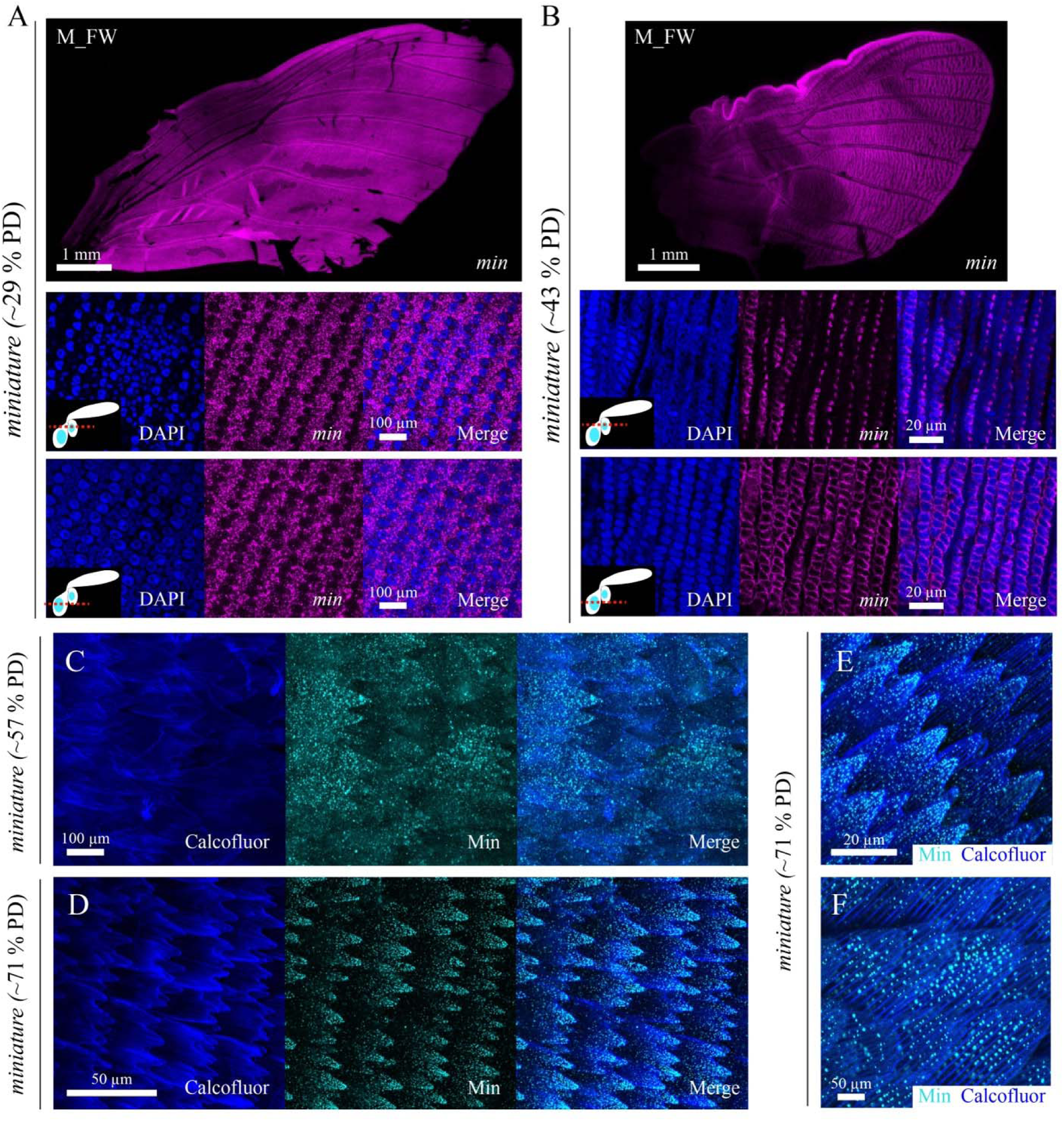
Miniature is dynamically expressed across the wing in all cuticle producing cell types. (A) At ~30% PD, *min* mRNA is expressed in all epithelial cells that make the bulk of the wing tissue. Below are two magnified views of a wing region at two different Z-planes marked by the insets. (B) At ~43% PD, *min* mRNA is no longer expressed in the epithelial cells of the wing but turns on in scale cells and is higher in the non-iridescent black regions. Below are magnified views of a wing region at two different Z-planes marked by the insets. (C) At ~57% PD, Min proteins appear in a speckled, punctate pattern across the developing scale blades. (D-F) At a later developmental timepoint, Min proteins are refined into clear puncta between the developing ridges and is concentrated at the finger-like projections of the scale tip.

CRISPR-Cas9 knockout of *min* dramatically affected both wing and scale development. Lethality was high but a few individuals emerged, smaller and with very crinkled wings in some cases (Supp Fig 15). Though *min* expression appeared to be higher in the black scales, scales all over the wing were affected. Many scales were tiny and cylindrical instead of flat (Supp Fig 15). Scanning electron micrographs (SEM) of these cylindrical scales showed unusual scale development (Fig 7A, Supp Fig 15). All cylindrical scales had a constriction just below the tip leading to a head of finger-like tentacles (Fig 7A, C). Over the scale body, pockets of cuticle bulged up and appeared to have sclerotized and hardened before being correctly structured (Fig 7B, D). In fact, various stages of development appeared to be ‘fossilized’ with some pockets unstructured (Fig 7B) and others taking the shape of nascent ridges (Fig 7D, Supp Fig 15). For scales that seemed to have developed normally and flattened, dentate scale tips were not formed and the ends of ridges stopped abruptly (Fig 7E).

**Figure 7:**
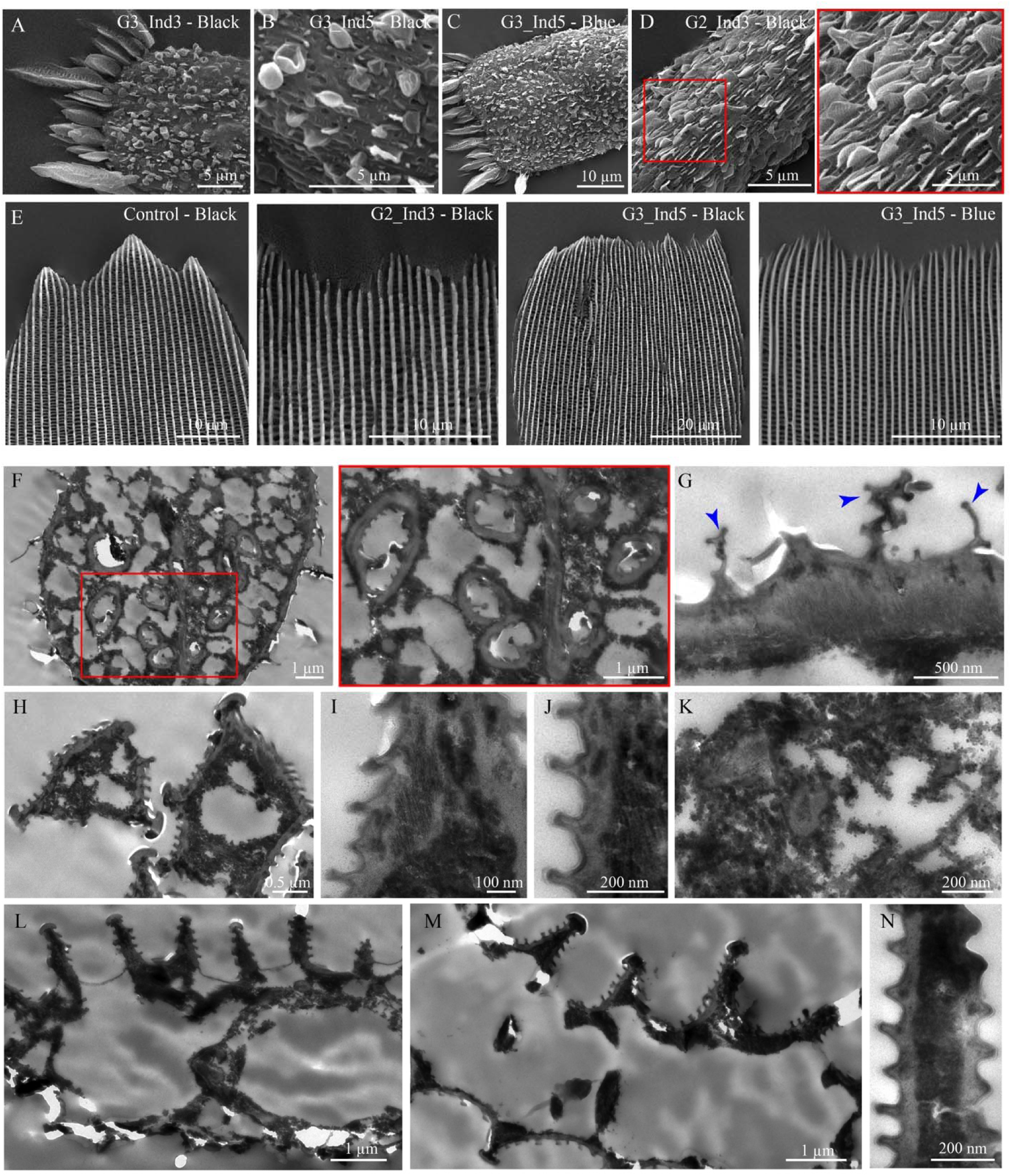
Miniature is necessary for correct scale cuticle formation and the development of dentate scale tips. (A-D) Scanning electron micrographs of various incorrectly formed scales in *min* crispant individuals. In strongly affected crispant scales which were cylindrical instead of flat, there was a consistent constriction just below the scale tip leading to a bunch of finger-like tentacles. Numerous pockets of cuticle dot the scale blade, potentially nascent ridges that buckled upwards but failed to develop fully across the scale before being sclerotized and ‘set’ by hardening. Red boxed area is magnified. (E) A control black scale with properly formed finger-like projections compared with flattened and expanded *min* crispant scales that lack ordered finger-like tip. (F) Transmission electron micrograph (TEM) cross-section of a cylindrical *min* crispant scale and a magnified view of the red boxed area. Numerous internalized bubbles of multi-layered cuticle with small ridges are seen. An electron-dense web-like mass of material has hardened within the scale body. (G) *min* crispant scale TEM cross-section with a clear electron-lucent layer buckled improperly, sitting atop a more electron-dense layer. Blue arrowheads indicate thin unstructured cuticle, possibly the envelope and epicuticle. (H-K) TEM cross-sections of the constricted finger-like tips and magnified images of the ridges and internal mass. (L-N) TEM cross-sections of flattened scales and a magnified view of a ridge.

Transmission electron micrograph (TEM) cross-sections provided an internal view of cuticle mis-formation. TEM images are from blue and black scales since it was hard to distinguish the scale types during sample preparation. Strikingly, in the small, cylindrical scales, bubbles of mini-cuticles with ridges were visible within the scale lumen, connected by a web-like network of electron-dense matter (Fig 7F). It appeared as if the scale collapsed on itself during development with many of the upper lamina nanostructures now forming within the collapsed scale. Closer inspection of the cuticle showed a thick electron-lucent layer with embedded regions of electron-dense matter, stacked on more electron-dense matter (Fig 7G). This was more clearly visible in the cross-sections of the finger-like tentacles (Fig 7H-K). Even in scales that were flattened and visually appeared to have developed correctly, TEM cross-sections showed a lack of proper integration of the cuticle layers and hazy masses of electron-dense matter within the scales and trabeculae (Fig 7L-N). The pockets and blobs of unstructured cuticle seen in the SEM images corresponded to only thin layers of cuticle, possibly the envelope and epicuticle (Fig 7G, blue arrowheads).

## Discussion

### Cuticle secretion and maturation dynamics affects nanostructure formation

In this study, we demonstrate key differences in the formation of the ECM determining structural differences in cell morphology. Comparing two scale types with different nanostructures, we showed that the initial stages of scale cuticle development i.e., the secretion of the first layer of the cuticle, the envelope, was initially similar between the blue and black scales. However, once the ridges were established by a buckling of the cell membrane (and the surrounding envelope) due to cell and envelope growth and the constrains provided by the underlying actin bundles (Totz et al. 2024; McDougal et al. 2021), structural differences in the envelope were seen between the blue and black scales. The blue iridescent scales exhibited a smooth envelope while the black scales developed a wavy envelope (Fig 1B, 50% PD). These early differences in the shapes of the envelope layer were likely instrumental in further divergence of ridge nanostructures as more of the epicuticle formed beneath the envelope. Blue scales developed a parallel lamella under the top of the ridge, which is important for the blue structural color, while the black scales developed multiple, smaller buckling of the envelope (Fig 1B, 64% PD).

What causes these early differences in envelope shape remains unknown but possible parameters include differences in the envelope composition between scale types, differential rate of envelope growth, rate of cuticle precursor secretion that forms the epicuticle and the composition of the cuticle precursors which will affect their maturation dynamics and hence the material properties of the secreted cuticle (Prakash et al. 2026).Wavy envelopes have been seen during nanopore development in *Drosophila* olfactory sensilla with the gene *Osiris23/gore-tex* playing an important role (Ando et al. 2019). *Osi* genes were identified in our omics studies and could be future targets to investigate. Gene knockouts only in certain regions of butterfly wings, to test questions of cuticle composition in specific scale types, remains challenging. However, mechanical probing methods such as Atomic Force Microscopy present future routes to test hypotheses of differences in cuticle properties between scale types such as stiffness (Prakash et al. 2026). We believe that rates of cuticle secretion and differences in its maturation, dependent on its material composition, are key factors that guide variations in nanostructures on butterfly scales (Prakash et al. 2026). In this context, it is an interesting question to consider the cellular mechanisms that determine rates of biomaterial secretion and the genes that might be involved in this regulation.

### The role of cuticle proteins and cuticle composition in butterfly scale development

Insect cuticles are complex structures that are composed of a mixture of biopolymers including multiple cuticle proteins (CPs). Over 200 CPs have been identified in insect cuticles and have been classified into 13 families (Willis 2010; Futahashi et al. 2008; Thakur et al. 2025). CP expression varies temporally and spatially during insect development and certain sub-families of CPs have been associated with certain types of cuticles like soft vs hard cuticles (Thakur et al. 2025).

In this study, we found many CPs to be differentially expressed between the blue and black tissues. The bulk nature of our omics data means that some of these differences will be due to tissue differences between the proximal (blue) and distal (black) regions of the wing, rather than differences between scale cell types per se. For example, large and thick trachea enter the proximal wing region where the wings are connected to the thorax and become thinner towards the distal wing tip. This likely explains why two of our candidate CPs, *Cpr100a* and *Cpr129*, that were enriched in the blue tissues in the omics data were associated with tracheal cuticle. In addition, since butterfly wings have both dorsal and ventral scales, as well as other cell types like epithelial and tracheal cells, we inevitably captured many of these cell types. Though the ventral scales for both the blue and black tissue regions under study were of a comparable color, i.e., brown, the similarities in the nanostructures of brown scales to the investigated black scales may have confounded our comparative analysis and the identified differentially expressed genes. In future, single-cell omics studies followed by bioinformatic separation of dorsal vs. ventral scale cells will help in the analysis of specific scale types.

*Cpr128* was strongly expressed in the distal wing regions corresponding to the black scales as predicted. However, this expression was localized to the socket cells at 57% PD. Though one crispant showed strong effects on scale development, we couldn’t detect any socket deformations. The mismatch between *Cpr128* mRNA localization and its functional role was puzzling. Perhaps the loss of one cuticular protein is complemented by other similar CPs during socket cuticle development. It is also possible that *Cpr128* mRNA is expressed in scale cells at later developmental time points which we couldn’t detect using a fluorescent based in situ hybridization, explaining the scale defects observed. Development of antibodies could help investigate protein expressions at later time points.

Our study also involved the functional verification of various candidate genes involved in scale nanostructure development using CRISPR-Cas9 targeted gene editing. Identification of targeted wing regions and mosaic patches were often challenging because scale nanostructure defects were not easily visible in some cases, unless entire wings were scanned using electron microscopy. Therefore, we might have missed many scale defects such as in our *Cpr128* crispants. Future experiments could attempt to create a transgenic butterfly line that expresses a fluorescent marker in all wing cells followed by co-injection of a double guide RNA (targeting the gene of interest and the fluorescent marker). Clonal patches lacking fluorescence would also have affected scales, making it easier to identify.

Nonetheless, we have shown that CPs are expressed in butterfly scale cells and play a role in scale development, like in moth scales (Liu et al. 2021). Combinations of different CPs expressed in different scale cell types will affect the material properties of the secreted cuticle such as its bending stiffness, in turn determining nanostructure formation. For example, in the beetle *Tribolium castaneum*, TcCPR27, TcCPR18 and TcCPR4 are abundant cuticle proteins in the rigid elytral procuticles. Knockout experiments have shown that these three CPs are important for the laminar architecture of the cuticle and the correct structure of the pore canal fibers (Noh et al. 2016, 2014). Correct localization of TcCPR27 is also necessary for the correct localization of TcCPR4 within the pore canals (Noh et al. 2015). Likewise, in the silkworm *Bombyx mori*, loss of BmorCPR2 changed the shape of the larvae because of a decrease in chitin binding, affecting the tensile properties of the cuticle (Qiao et al. 2014). Characterizing highly expressed CPs in different scale cell types will therefore be crucial in understanding how cells influence the biomechanics of their secreted ECMs.

### Zona pellucida domain proteins are important ECM components of butterfly scales

A characteristic motif found in many ECM proteins is the ~260 amino acid Zona pellucida domain. Members of the ZP family of proteins are expressed in cuticle secreting cells, enriched in the apical ECM and have self-assembly properties guided by the polymerization potential of the ZP domain, leading to the formation of crosslinked protein fibrils and matrices (Plaza et al. 2010). Maturation of ZP proteins is a complex process involving numerous post-translational modifications including glycosylation. Some ZP proteins are anchored to the plasma membrane and bridge the membrane to the apical ECM while others are cleaved and secreted into the extracellular space forming scaffolds or as part of the cuticle (Ray et al. 2015; Drees et al. 2023).

The role of ZP proteins in controlling cell and tissue shapes have been well documented with many studies highlighting their important roles in governing epithelial nanostructures across species (Cohen et al. 2020; Su et al. 2026; Zhang et al. 2023). In *Drosophila melanogaster*, numerous examples exist. In embryonic denticle development, different ZP proteins are organized in specific sub-regions of the apical compartment and mediate the link between the ECM and cell membrane (Fernandes et al. 2010). Disruptions of these ZP proteins affects denticle shapes in its corresponding apical sub-compartment (Fernandes et al. 2010). Similarly, the characteristic shapes of *Drosophila* appendages like the wings are determined by opposing biomechanical forces generated by cell constriction on one hand vs localized anchorage to the ECM on the other hand, the latter mediated by ZP proteins like Dumpy (Ray et al. 2015). More recently, Dusky-like (Dyl) has been implicated in correct biconvex corneal lens development (Ghosh and Treisman 2024). Not only does it stabilize the expansion of the apical cell membrane, it also determines the correct positioning of other ZP proteins like Dumpy and Piopio external to the corneal lens, creating a scaffold within which chitin and other biomolecules secreted by the underlying cells form the lens (Ghosh and Treisman 2024). These external protein scaffolds are then likely degraded by proteolysis (Ghosh and Treisman 2024). In bristles, Dyl is needed for correct chitin deposition (Nagaraj and Adler 2012). Similarly, in the olfactory sensilla, ZP proteins are necessary for correct envelope waviness and formation of nanopores that permit odorant molecule entry (Itakura et al. 2026). Due to differential affinities of ZP proteins, they form different layers before cuticle secretion, determining the shape of the envelope. Dyl is important for the formation of the envelope which develops at the border between different ZP protein layers. Other ZP proteins like Tyn, Nyobe, Neo and Morpheyus form a cloud ECM around the sensilla providing a compressive mechanical force restricting sensilla growth and likely determining correct envelope shape (Itakura et al. 2026). We didn’t observe a cloud ECM around the scales but there was a membrane-proximal zone above the plasma membrane, at the edge of which the envelope developed (Fig 1B, 43% PD).

We identified similar functions for ZP proteins in determining butterfly scale shapes and nanostructure formation. Like in *Drosophila* denticles and olfactory sensilla, we saw some sub-compartmentalization of the apical membrane along the scale’s long axis, with Tyn being strongly expressed at the base where the scale cell exits the socket (Fig 5). Mutations of *tyn* strongly affected that sub-compartment with scales losing the bulge in their stalk and having pointy “tails”. *tyn* crispants also displayed curved and malformed scales with mosaic patches of ridges without lamellae (or distantly spaced lamella) and covered inter-ridge spaces without windows. The *min* crispants showed the strongest phenotypes with butterflies eclosing with small and crumpled wings. This phenotype was expected because *min* was expressed in epidermal cells across the entire wing during early pupal development in *H. sara* and is known for its role in *Drosophila* wing morphogenesis (Roch et al. 2003). The strong scale mutations however were very interesting and suggests that Min plays additional important roles in the formation of the initial scale ECM. First, *min* crispant scales that flattened did not display dentate edges with finger-like projections implicating *min* in finger formation. This was in line with concentrated Min protein expression in the scale fingers. Second, the small cylindrical scales with blobs of unstructured cuticle or partially developed ridges with primitive ridge structures suggests that the structural integrity of scales was lost in *min* crispants, likely due to abnormal cuticle formation. It is likely that Min is a critical component of the epicuticle or is necessary to anchor the developing cuticle to the plasma membrane at critical locations. Without Min protein, formation of a crosslinked epicuticular matrix may have been structurally unsound or lacking points of attachment. With different biomechanical forces acting within and outside the cell as the cell grows, a weak ECM may lead to restructuring of the scale surface into a cylinder instead of a paddle to account for a stable configuration. Perhaps Min interacts with other ZP proteins such as Tyn and organizes into different layers. The lack of integration of the different cuticle layers and the appearance of electron dense regions within electron lucent layers of the *min* adult scales, as compared to a homogenous electron dense mass in wildtype scales, strengthens our conclusions that Min is an integral ECM component.

Our work provides the first evidence that ZP proteins are involved in controlling the complex morphology of butterfly scales, and more broadly adds to the growing body of evidence that they are involved in mechanochemical processes controlling cell and ECM shape generally. Further investigation of ECM-associated ZP proteins in a wider range of systems and organisms could reveal more about the general processes governing the formation of complex cell shapes. This work also provides a starting point for greater understanding of optical nanostructure formation specifically in butterfly scales. Our results present a plethora of exciting future avenues to investigate. Are Tyn and Min secreted outside the cell into the extracellular space or do they act as attachment points between the membrane and the ECM via their transmembrane domains? Why do the tiny cylindrical scales in *min* crispants always have a constriction below the scale tip leading to a “medusa” scale? How is the sub-compartmentalization of the scale apical ECM determined? Are other ZP proteins, not investigated in this study, such as Dyl, Nyobe, Neo, Morpheyus or Dumpy, important ECM components in butterfly scales? Using butterfly scales and their diversity of nanostructures as a model, our study identifies some of the downstream members of the genetic networks that regulate scale development, such as cuticle proteins and ECM components, presenting a foundation for future explorations on this topic. Information derived from techniques such as single-cell analyses, super-resolution microscopy and mathematical modelling can help improve our understanding of the development of biological nanostructures.

## Materials and methods

### Butterflies

*Heliconius sara* butterflies were reared in temperature and humidity-controlled chambers at the Arthur Willis Environment Centre, University of Sheffield. Adults were fed with a sugar solution supplanted with pollen and eggs were collected on stems of the host plant *Passiflora auriculata*. Caterpillars were reared in tanks on *Passiflora biflora* until pupation. Pupation time was noted, and it lasted 8-9 days in this species at 24°C and ~7 days at 26-27°C.

### Sample embedding and transmission electron microscopy

Small wing pieces from the blue and black regions of the forewing at various developmental time points were dissected using a blade. Samples were fixed overnight at 4°C in 2.5% glutaraldehyde in 0.1M sodium cacodylate buffer followed by three washes with 0.1M sodium cacodylate. Postfixation was carried out in 2% Osmium tetraoxide in deionized water for 1 hour at room temperature under a fume hood and then washed in deionized water three times. Dehydration was through an ascending ethanol series at room temperature: 25% ethanol for 5 minutes, 50%, 75%, 95% and 100% ethanol for 20 minutes. Samples were then transferred to 100% propylene oxide for 10 minutes, 2 times. Infiltration was with the following steps: a) 100% propylene oxide: Araldite resin (3:1) for 30 mins at room temperature b) 100% propylene oxide: Araldite resin (1:1) for 1 hour at room temperature c) Leave tubes open for 30 minutes under a fume hood for the propylene oxide to evaporate d) Pure Araldite resin overnight at room temperature d) Samples moved to blocks with pure Araldite resin for 1 hour in a 40°C oven e) Blocks moved to a 50°C oven for 1 hour f) Finally, blocks were moved to a 60°C oven. Samples were reoriented constantly until the resin was sufficiently hardened to prevent samples from moving. Resin blocks were allowed to polymerize at 60°C for 1-2 days.

Ultrathin sections were cut using a Reichert-Jung Ultracut E Ultramicrotome and transferred to either formvar coated or carbon coated grids. Grids were stained with 2% aqueous uranyl acetate for 30 minutes in the dark and washed in deionized water for 30 minutes. Grids were then stained with lead citrate for 5 minutes followed by another deionized water wash for 5 minutes. Images were acquired on a FEI Technai T12 Sprit Transmission Electron Microscopy (FEI, Thermo Fisher Scientific, U.S.A).

### Tissue dissection for RNA extraction and sequencing

Samples for the RNA-seq experiment were taken from across six separate batches of caterpillars between December 2021 and May 2022. Butterflies were reared at 24°C with the pupation period lasting roughly 9 days. Wing pieces were collected at five pupal time points – Day 2 (22%), Day 3 (33%), Day 4 (44%), Day 5 (56%) and Day 6 (67%). At each time point, five biological replicates were collected with one individual per replicate (both forewing pieces pooled) except 22% and 33% development, in which two individuals were pooled at this stage to make up a single sample. Both forewings were dissected in Graces Insect Medium (room temperature) with presumptive blue and black regions of the forewings were cut based on vein markings and immediately placed in tubes of RNAlater at 4. After several hours tubes were transferred to −80 for storage until RNA extraction.

RNA was extracted with Qiagen RNAeasy kit with residual DNA removed using on-column DNase digestion using the RNAse-Free DNase Set. Library preparation, sequencing and quality control was performed by the NERC Environmental Omics Facility (NEOF), University of Liverpool. A dual-indexed, RNAseq library was prepared from the total RNA using NEBNext Poly(A) selection and UltraII Directional RNA library preparation kits. Sequencing was performed on a single lane with an Illumina NovaSeq S4 (paired end, 2x 150bp). RNA-seq data have been deposited at GEO with the project accession number GSE339405.

### RNA-seq quality control, read alignment, transcript assembly and quantification

Quality control was performed on the raw fastq files by trimming off the illumina adapter sequences using Cutadapt (v. 1.2.1) with the option -O 3. The reads were further trimmed using Sickle (v. 1.200) with a minimum window quality score of 20. Reads shorter than 15 bp after trimming were removed. Reads in which only one of the pair passed the filtering stage were assigned to separate files (and not subsequently used).

All alignment and assembly steps were performed on the University of Sheffield High Performance Computing (HPC) cluster, ShARC. Filtered reads were downloaded from the sequencing facility in fastq format. Read quality was visualised using FastQC (version 0.11.8) and the resulting quality control files aggregated and inspected using multiQC (version 1.5). Using HISAT2 (version 2.1.0) (Kim et al. 2015)the trimmed reads were aligned to a reference *Heliconius sara* genome downloaded from NCBI (GenBank assembly accession: GCA_917862395.1) https://www.ncbi.nlm.nih.gov/datasets/genome/GCA_917862395.1/.

HISAT2 was performed with the ‘--knownsplicesite-infile’ option to enable alignment of small anchor reads. Exon and splice site information was extracted from the gene annotation file (Hsar.v1.1.annotation.CAT.gff3) provided by (Cicconardi et al. 2023). Gffread was used to convert the gene annotation file into ‘.gtf’ format and ‘Hisat2_extract_splice_sites.py’ was used to create a list of splice sites for HISAT2. In addition, HISAT2 was run with the -dta option to adapt the alignments for downstream use in StringTie. Output SAM files from HISAT2 were sorted and converted into .BAM files using SAMtools (version 1.9) (Li et al. 2009).

StringTie (v 2.1.5) was used to perform transcript assembly and read quantification (Pertea et al. 2015). The gene annotation file was provided as an input to guide read assembly and gtf files generated for each sample. StringTie was then run in ‘merge mode’ to create a uniform set of transcripts (Stringtie_merged.gtf). This non-redundant transcript set was then used to re-estimate abundance of the output alignment files from HISAT2 using the ‘-e’ option. Output Table (ctab) files provided coverage data. The R package ‘IsoformSwitchAnalyzeR’ (v. 1.20.0) was used to assign gene names to the transcripts assembled by StringTie. The ImportIsoformExpression() and importRdata() functions were used to import the ctab files generated by StringTie and gene annotation file into R and the isoform annotations recovered. The extractGeneExpression() function was used to generate a read count matrix for downstream differential expression analysis in EdgeR.

### Sex determination in the RNA-seq data

Sex of the pupae was not identified prior to sequencing, so we used the sequence data to determine the sex of the individuals based on heterozygosity of the Z (sex) chromosome (Midic et al. 2018). SNP calling was performed on the BAM files with GATK HaplotypeCaller (v. 4.1.0.0) (Poplin et al. 2018; McKenna et al. 2010). Alignments and bases with a mapping quality <20 were removed. A custom Perl script was used to count the number of heterozygous and homozygous sites and calculate the proportion of heterozygous sites. As females in *Heliconius* are the heterogametic sex (ZW) (Sahara et al. 2012), we expect values of heterozygosity for Z chromosome SNPs to be zero, although a relatively minor amount of heterozygosity was observed due to base calling errors. The proportion of heterozygous sites across all samples showed a bimodal distribution with all individuals either <0.05 or >0.15, so all samples above 0.15 were assigned as male. As individuals were pooled for sequencing at 22% and 33% development, we could not assign these samples to a sex, due to the possibility that they were mixed-sex. In the later stages, where sex could be assigned, there was no separation of the sexes in the PCA and so sex was not considered in subsequent analyses.

### Differential expression analysis

Differential expression analysis was conducted in R using the R/Bioconductor package EdgeR (v. 3. 40. 2) (Robinson et al. 2010). Genes with low expression values were removed from the gene count matrix using the filterByExpr() function (Chen et al. 2016). Normalisation of library size was then performed using the trimmed mean of M-values (TMM) method (Robinson and Oshlack 2010). Samples were clustered using Multidimensional Scaling to check for any anomalies. This revealed two individuals (four samples) sampled at day 4 and day 6 that clustered with different developmental timepoints from expected. This may be due to mislabelling of the pupation date or variation in developmental rates between individuals (some individuals took 8 rather than 9 days for pupation). To remove noise introduced by this variation, we removed these samples from further analysis. A further pair of samples from day 2 showed opposite clustering of the blue and black tissues compared to all other individuals and was therefore likely mislabelled. Further inspection of expression values using heatmaps supported the mislabelling of the tissues as the expression values matched the other tissue. The metadata was corrected for the labelling mistake. In total 46 samples were used in the analysis.

### Wing scales/cell dissociation and purification for proteomics

For the proteomics experiment, butterflies were reared at 24°C with the pupation period lasting roughly 8 days. Wing scales and cells were collected at four pupal time points – Day 4 (~50% of pupal development), Day 5 (~64%), Day 6 (~76%) and Day 7 (~88%). At each time point, five biological replicates were collected with one individual per replicate (2 forewings). All individuals sampled were males. Both forewings were dissected in ice cold 1X Phosphate Buffered Saline (PBS). Since Day 4 wings were fragile, forewings were dissected with the pupal case intact. Presumptive blue and black regions of the forewings were cut based on vein markings and transferred into well plates with 1 mL cold 1X PBS. Cut wing tissues were crushed with a fine brush until completely dissociated. For the fragile Day 4 wing pieces, trituration with a pipette tip was enough to mechanically dissociate the tissues. Day 4 and Day 5 samples were then filtered through a 40 µm filter while Day 6 and Day 7 samples were filtered through a 100 µm filter into a new 1.5 mL Eppendorf tube. Samples were centrifuged at 12000 rpm, 10 minutes, 4°C to obtain a large pellet at the bottom that contained dissociated cells and individual scales. PBS was decanted as much as possible, and the pellets were stored at −80°C.

### Protein extraction and quantification

Proteins were extracted from all the sample pellets using a lysis buffer containing 5% SDS and 50 mM Triethylammonium bicarbonate (TEAB) pH 8.0. 40 µl of lysis buffer was added to each sample and heated at 90°C for 15-20 minutes, shaking at 1200 rpm. Samples were then centrifuged at 20000g for 10 minutes. 35 µl of the supernatant was transferred to new Eppendorf tubes without disturbing any pellets if present. Protein concentrations were measured using a Micro BCA Protein Assay Kit (Thermo Fischer Scientific, Catalog #23235) with 2.5 µl of extracted protein from each sample. Samples were then assigned random numbers to anonymize the downstream proteomics analyses.

### Protein reduction, alkylation and trypsin digestion using S-traps

All samples were diluted as needed with lysis buffer to obtain 20 µg of starting protein. Samples were reduced by adding 1:10 volume of 220 mM of Dithiothreitol (final concentration of 20 mM) and incubated at 95 °C for 10 minutes at 800 rpm. Samples were brought to room temperature and alkylated by adding 1:10 volume of 440 mM Iodoacetamide (final concentration of 40 mM) and kept in the dark at room temperature for 30 minutes without shaking. 1:10 volume of 25% phosphoric acid was added (final concentration of 2.5%) followed by 7 times the volume of binding buffer (1:10 dilution of 1M TEAB pH 7 in methanol). Samples were then digested using S-trap technique according to manufacturer’s protocol (Protify, USA). In short, samples were loaded in batches into S-trap columns and centrifuged at 4000g for 1 minute. S-traps were washed with 150 µl of binding buffer five times, discarding flow-through to waste as necessary. A final spin at 4000 g, 1 minute was carried out to remove all the wash buffer. 1:10 amount of trypsin to the starting amount of protein i.e., 2 µg of trypsin, was added to the S-trap making sure there was no bubble between the column and trypsin. The trypsin was reconstituted in 50 mM TEAB and a minimum volume of 25 µl was added on top of the column. A cut end of a 1 mL pipette tip was used to apply pressure to the top of the S-trap column until the trypsin passed through the column. The S-trap columns were loosely capped, transferred to a clean 1.5 mL Eppendorf tube, sealed with parafilm and incubated at 37 °C overnight with no shaking.

Tubes were centrifuged quickly after overnight incubation, then eluted with three buffers. 40 µl of Elution 1 (50 mM TEAB) was added to each tube and centrifuged at 4000g, 1 minute. 40 µl of Elution 2 (0.2% formic acid) was then added followed by another centrifugation step. Finally, 35 µl of Elution 3 (50% acetonitrile with 0.2% formic acid) was added to each tube and centrifuged at 4000 g, 1 minute. Samples were then dried in a vacuum concentrator.

### LC-MS/MS Analysis

Proteomic analyses were performed at the biOMICS Mass Spectrometry Facility, University of Sheffield, using an Orbitrap Exploris 480 mass spectrometer (Thermo Fisher Scientific) equipped with a nanospray ionisation (NSI) source and coupled to a Vanquish HPLC system (Thermo Fisher Scientific). Peptides were desalted online using a nano-trap column (75 μm i.d. × 20 mm; Thermo Fisher Scientific) and subsequently separated using an EASY-Spray column (50 cm × 50 μm i.d., PepMap C18, 2 μm particle size, 100 Å pore size; Thermo Fisher Scientific). Peptide separation was performed using a 100-min gradient with buffer B consisting of 0.5% (v/v) formic acid in 80% (v/v) acetonitrile. The gradient increased from 3% to 20% buffer B over 68 min, followed by an increase to 35% buffer B over 23 min and then to 99% buffer B over 1 min. The column was subsequently maintained at 99% buffer B for 9 min.

The mass spectrometer was operated in positive-ion mode using a data-dependent acquisition (DDA) method. The nanospray voltage was set to 1.9 kV and the ion-transfer-tube temperature to 275°C. Full MS spectra were acquired in the Orbitrap over an *m/z* range of 375–1200 at a resolution of 120,000, with a normalized automatic gain control (AGC) target of 300% (absolute AGC target, 3 × 10), automatic maximum injection time and one microscan.

For data-dependent MS/MS acquisition, precursor ions exceeding an intensity threshold of 1 × 10 and with charge states of 2–5 were selected. Up to 20 precursor ions were selected for fragmentation per acquisition cycle. Dynamic exclusion was set to 45 s after a single selection, with a ±10 ppm mass tolerance, and isotope exclusion was enabled. Precursors were isolated using a 2 *m/z* isolation window and fragmented by higher-energy collisional dissociation (HCD) at a normalized collision energy of 30%. MS/MS spectra were acquired in the Orbitrap at a resolution of 15,000, with the AGC target set to the instrument standard setting, automatic maximum injection time and one microscan. MS/MS spectra were acquired in centroid mode with the first mass set to *m/z* 120.

### MS Data Processing and Protein Identification

Raw mass spectrometry data were processed using MaxQuant (version 1.6.10.43). MS/MS spectra were searched against a custom protein sequence database generated from a translation of the RNAseq data and contained 114,380 sequences. Database searches were performed using Trypsin/P as the proteolytic enzyme, allowing up to two missed cleavages. Carbamidomethylation of cysteine residues was specified as a fixed modification, whereas oxidation of methionine and protein N-terminal acetylation were specified as variable modifications.

Peptide and protein identifications were controlled at a false discovery rate (FDR) of 1% using the target-decoy strategy implemented in MaxQuant. FDR thresholds of 0.01 were applied at both the peptide and protein levels. Label-free protein quantification was performed using the MaxQuant label-free quantification (LFQ) workflow.

### Label-Free Quantitative Proteomics and Statistical Analysis

The MaxQuant output was imported into Perseus (version 1.5.6.0), and LFQ intensity values were used for quantitative analysis. Protein entries corresponding to potential contaminants and reverse-database matches were removed prior to statistical analysis.

Proteins were retained for further analysis when valid LFQ intensity values were detected in at least 70% of samples in at least one experimental group. LFQ intensity values were log - transformed prior to statistical analysis. Log - transformed intensities were subsequently normalized to the median, and missing values were imputed from a normal distribution.

Differential protein abundance between blue and black tissue groups was assessed using a two-sample *t*-test with a permutation-based FDR of 0.05. Proteins exhibiting a fold change >1.5 together with a *p*-value <0.01 were considered significantly differentially abundant and were retained for subsequent analyses.

### Proteomics Data Availability

The mass spectrometry proteomics data have been deposited to the ProteomeXchange Consortium via the PRIDE (Perez-Riverol et al. 2025) partner repository with the dataset identifier PXD080755

### SDS-PAGE Gel

Three samples from each time point were run on PAGE gels with a 4 % stacking gel and 12 % resolving gel (SureCast Gel Handcast System, Invitrogen). 10 µl of each sample was mixed with 10 µl of 2X loading dye at 95°C for 10 minutes. All 20 µl of the samples were loaded on the gels and run at 140V for 1 hour along with 5 µl of PageRuler protein ladder (Thermo Scientific; Cat #26616). Gels were fixed in Fix solution (40% Methanol and 2% Acetic Acid) for 30 minutes followed by two rinses with water. Gels were then stained overnight in diluted Brilliant blue solution and destained the following day with 20% methanol.

### Functional annotation and gene set enrichment analysis

All *H. sara* genes in the transcriptomic gene count matrix were used for functional gene annotation (n = 16901) by identifying homology to *Drosophila* genes. Fasta sequences of the genes were extracted from the reference NCBI *H. sara* genome using Gffread and the StringTie merged gtf file. Using the ‘makeblastdb’ function from ncbi-blast (version 2.8.1) a blast database was created for all *Drosophila* protein sequences downloaded from UniProtKB (https://www.uniprot.org/uniprotkb?query=drome&facets=model_organism%3A7227). The *Drosophila* database was then interrogated for the *H. sara* transcript sequences using ‘blastx’ with default parameters. Hits were then sorted and scored based on length and E-value to select the best hit for each *H. sara* gene and hits with E-value > 1×10^−5^ discarded. In total, 11,068 *H. sara* genes showed a good match to *Drosophila*.

Gene set enrichment analysis was conducted using PANGEA (version 2) (Hu et al. 2023). For the transcriptomic analysis we used all annotated genes from the gene count matrix as the background gene list, and for the proteomic analysis we used all annotated proteins found in the proteomic data, using the gene symbols based on homology to *Drosophila*.. These background sets were compared to the significantly DE genes and proteins (FDR<0.05) separately at each developmental stage, using Direct Gene Ontologies (GO) gene sets for Molecular Function, Biological Process and Cellular Components. GO terms with Bonferroni corrected P-values less than 0.05 were classed as significantly enriched.

### Immunostainings

A rabbit anti-miniature and rat anti-trynity primary antibodies were kind gifts from Dr. Hélène Chanut at the University Paul Sabatier Toulouse – CNRS. The rabbit anti-ds primary antibody was a kind gift from Dr. David Strutt, University of Sheffield, UK. All three antibodies were raised against *Drosophila* antigens. Alexa Fluor 488 goat anti-rabbit IgG (Thermo Fisher Scientific Cat #A-11034), Alexa Fluor 568 goat anti-rat (Thermo Fisher Scientific Cat #A-11077) and an Alexa Fluor 555 goat anti-rabbit IgG (Thermo Fisher Scientific Cat #A-21428) were used as secondary antibodies. To stain for actin, either an Alexa Fluor 555 Phalloidin or ATTO-647 Phalloidin (Invitrogen) was used.

Pupal wing tissues were dissected at designated time points and fixed in 4% paraformaldehyde in PBS at room temperature for 20-30 minutes. Wings were then washed three times in PBS and incubated in block buffer (PBS + 0.1% Tween-20 + 1% BSA) for 2 hours at room temperature. Wings were then incubated in primary antibody (1:500 for anti-miniature and anti-trynity; 1:50 for anti-ds) in wash buffer (PBS + 0.01% Tween-20 + 0.25% BSA) overnight at 4°C. Samples were washed 3-4 times with wash buffer followed by incubation with secondary antibodies (between 1:200 and 1:500) for 1 hour at room temperature. The wings were then incubated with DAPI (1:1500) for 30 minutes if required, and washed with wash buffer again for 4 times, 20 minutes each. Wings were mounted on glass slides using clarified Mowiol 4-88 (Sigma Aldrich, Cat#81381) and imaged either on a Nikon A1 Confocal (Nikon, Japan) or a Zeiss Airyscan (Zeiss, Germany).

### Hybridization chain reaction (HCR3.0 – Fluorescent based in situ hybridization)

Fluorescent *in situ* hybridization of various mRNA targets followed the protocol described in Choi et al., 2018 (Choi et al. 2018)with a few modifications in buffers and incubation conditions. Timed pupal wings were fixed in 4% formaldehyde in PBS at room temperature for 20-30 minutes. Wings were thereafter washed thrice with 1x PBST and treated with a detergent solution followed by three washes with 1x PBST and two washes with 5x SSCT. Wings were then transferred to a hybridization buffer. Hybridization involved incubation in a solution containing 20 mL (100 mM) of probe set against each gene (Sigma Aldrich) in 1 mL of probe hybridization buffer at 37°C overnight followed by four washes with probe wash buffer at 37°C, 20 minutes each. Afterward, wings were washed with 5X SSCT and incubated in an Amplification buffer for 30 minutes. For the chain reaction, a solution with HCR hairpins (B1-Alexa Fluor 647; Molecular instruments) in the amplification buffer was added to the tissues and incubated in the dark overnight. Washes were carried out with 5x SSCT and counterstaining was with DAPI (1:1500) for 30 minutes at room temperature. Wings were mounted on glass slides using clarified Mowiol 4-88 (Sigma Aldrich, Cat#81381) and imaged on a Nikon A1 Confocal (Nikon, Japan). Images were post-processed using Fiji to improve contrast. The primers for HCR are specified in Supplementary Table 3 (excel file) and the composition of the various HCR buffers is provided in Supplementary Table 4.

### CRISPR-Cas9 gene editing

CRISPR-Cas9 targeted gene editing was performed using the following protocol. Eggs were collected for 1-1.5 hours on a shoot of *Passiflora auriculata.* Since *Heliconius sara* females lay eggs in big clumps, eggs clumps were dissociated by placing them in a 1:30 Milton’s solution for 6 minutes with gentle shaking. Eggs were then washed twice with water and any undissociated eggs were gently broken apart with a brush. Individual eggs were arranged in rows on double-sided sticky tape.

Guide DNA templates against regions of interest in target genes (Supplementary Table 5) were manually designed following the GGN_18_NGG pattern to produce forward primers 5’-GAAATTAATACGACTCACTATAGG-xxxxxxxxxxxxxxxxxxxxx-GTTTTAGAGCTAGAAATAGC-3’. Double stranded DNA templates were generated using PCR with Q5 High-Fidelity DNA polymerase (NEB Cat # M0491S) and a common reverse primer for all targets 5’-AAAAGCACCGACTCGGTGCCACTTTTTCAAGTTGATAACGGACTAGCCTTATTTTAA CTTG CTATTTCTAGCTCTAAAAC-3’. Purified DNA templates were used for *in vitro* transcription reactions with a T7 RNA Polymerase (NEB Cat # M0251S) to generate guide RNAs that were then purified using an ethanol precipitation. Injection mixtures containing different concentrations of Cas 9 (NEB Cat # M0641) protein and different guide RNAs (Supplementary Table 6) were injected into the eggs within 4-6 hours of egg laying. Hatched caterpillars were transferred onto a small shoot of *Passiflora biflora* in a petri dish for 1-2 days before being transferred into larger tanks. Adults were scored for their phenotypes.

### Genotyping crispants

Genomic DNA from the legs and thoracic tissues of emerged adults and pieces of preserved pupae from which butterflies didn’t eclose was extracted using the PCRBIO Rapid Extract PCR Kit (PCR Biosystems). Genomic DNA from controls (injected with just Cas 9 and no guide RNA or not injected at all) was also extracted. Targeted genes of interest were amplified using primers (Supplementary Table 5) from both the crispants and control, purified using ethanol purification and Sanger sequenced. Crispants were identified using the online Inference of CRISPR Edits (ICE) tool (Conant et al. 2022).

### Scanning electron microscopy

Individual scales were sampled by picking them using insect pins while entire wing pieces were cut using scissors. Samples were placed on double sided carbon tapes on SEM stubs and sputter coated with gold. Images were acquired on a Tescan Vega 3 Scanning Electron Microscope.

### Statistical analysis

Statistical analyses were performed in R Studio 2026.04.0+526 with R 4.5.2 (R Core Team 2021). The differences in mean bulge width between wildtype and *tyn* crispants were analyzed using a linear mixed-effects model (LME) via the nlme package (v 3.1.152) (Pinheiro et al. 2021). To account for potential non-independence of scales measured from the same butterfly, we evaluated random-effects structures including individual butterfly (Ind) as a random intercept and scales nested within individual (Ind/Scale). Models were evaluated using maximum likelihood (ML) estimation and compared via Akaike Information Criterion (AIC) and likelihood ratio tests (LRT). The nested random effect of scale (Ind/Scale) did not improve model fit over an individual-level random effect alone (LRT < 0.001, p = 0.999), nor did accounting for heteroscedasticity across sample types using a varIdent variance structure significantly improve fit (LRT = 3.00, p = 0.083). Therefore, the most parsimonious model—retaining individual butterfly (Ind) as a single random intercept—was selected and refitted using Restricted Maximum Likelihood (REML) for final inference. Three individuals were measured for wildtype (8-15 scales per individual) and two *tyn* crispants (15 scales each).

## Supporting information

Supplementary Table 3 - Primers for HCR

## Competing Interests

The authors declare no competing interests.

## Acknowledgements

We thank Jonathan Paul Richards for maintaining the *Heliconius sara* butterfly stock at the University of Sheffield. Dr. Chris Hill and Dr. Svetomir Tzokov of the Electron Microscopy Unit at the University of Sheffield gave advice and help in preparing the TEM resin blocks and sections. We are grateful to Laura Hartshorne and Dr. Mirre Simons for the use of their microinjector. We acknowledge the support of Lucy Knowles, Tom Holden, Gavin Horsburgh and Rachel Patel of the NEOF facility, University of Sheffield, for their lab support and useful discussions, and the NEOF facility staff at the University of Liverpool for preparing and sequencing the RNA-seq libraries. We are thankful to Dr. Hélène Chanut at the University Paul Sabatier Toulouse – CNRS for providing the Miniature and Trinity antibodies and Dr. David Strutt, University of Sheffield for the Dachsous antibody. Confocal imaging work was performed at the Wolfson Light Microscopy Facility, with training and support from Dr. Darren Robinson and Dr. Nick van Hateren. We are grateful to Dr. Mathias Kolle and Dr. Bodo Wilts for useful discussions on the topic of butterfly scale development and Dr. Ellen Allwood for help and discussions around proteins. This work was funded by a Human Frontier Science Program Grant (RGP0034/2021) and a Medical Research Council grant to MOC (MR/X012220/1).

## Author Contributions

A.P. and N.N. conceived the study. A.P., V.L., V.S. and R.G. performed the experiments. A.P., V.L., Y.B.C., R.G., T.K.P. and N.N. performed the formal analysis. A.P. wrote the original draft of the manuscript. A.P. and N.N. reviewed and edited the manuscript with contributions from all authors. M.O.C., and S.A. provided guidance and support. N.N. acquired the funding and supervised the study.

## Notes

### Competing Interest Statement

The authors have declared no competing interest.

## References

Ando T, Sekine S, Inagaki S, Misaki K, Badel L, Moriya H, Sami MM, Itakura Y, Chihara T, Kazama H, et al. 2019. Nanopore Formation in the Cuticle of an Insect Olfactory Sensillum. Current Biology 29: 1512–1520.e6. 10.1016/j.cub.2019.03.043.

Bosveld F, Bonnet I, Guirao B, Tlili S, Wang Z, Petitalot A, Marchand R, Bardet P-L, Marcq P, Graner F, et al. 2012. Mechanical Control of Morphogenesis by Fat/Dachsous/Four-Jointed Planar Cell Polarity Pathway. Science (1979) 336: 724–727. 10.1126/science.1221071.

Chen Y, Lun ATL, Smyth GK. 2016. From reads to genes to pathways: differential expression analysis of RNA-Seq experiments using Rsubread and the edgeR quasi-likelihood pipeline [version 2; peer review: 5 approved]. F1000Res 5.

Choi HMT, Schwarzkopf M, Fornace ME, Acharya A, Artavanis G, Stegmaier J, Cunha A, Pierce NA. 2018. Third-generation in situ hybridization chain reaction: multiplexed, quantitative, sensitive, versatile, robust. Development 145: dev165753. 10.1242/dev.165753.

Cicconardi F, Milanetti E, Pinheiro de Castro EC, Mazo-Vargas A, Van Belleghem SM, Ruggieri AA, Rastas P, Hanly J, Evans E, Jiggins CD, et al. 2023. Evolutionary dynamics of genome size and content during the adaptive radiation of Heliconiini butterflies. Nat Commun 14: 5620. 10.1038/s41467-023-41412-5.

Cohen JD, Bermudez JG, Good MC, Sundaram M V. 2020. A C. elegans Zona Pellucida domain protein functions via its ZPc domain. PLoS Genet 16: e1009188-. 10.1371/journal.pgen.1009188.

Conant David, Hsiau Tim, Rossi Nicholas, Oki Jennifer, Maures Travis, Waite Kelsey, Yang Joyce, Joshi Sahil, Kelso Reed, Holden Kevin, et al. 2022. Inference of CRISPR Edits from Sanger Trace Data. CRISPR J 5: 123–130. 10.1089/crispr.2021.0113.

Dinwiddie A, Null R, Pizzano M, Chuong L, Leigh Krup A, Ee Tan H, Patel NH. 2014. Dynamics of F-actin prefigure the structure of butterfly wing scales. Dev Biol 392: 404– 418.

Drees L, Schneider S, Riedel D, Schuh R, Behr M. 2023. The proteolysis of ZP proteins is essential to control cell membrane structure and integrity of developing tracheal tubes in Drosophila eds. E. Knust and C. Desplan. Elife 12: e91079. 10.7554/eLife.91079.

Duan Y, Merzendorfer H, Yang Q. 2026. Molecular Insights into the Biosynthesis of Insect Cuticles. 10.1146/annurev-ento-121423-.

Fernandes I, Chanut-Delalande H, Ferrer P, Latapie Y, Waltzer L, Affolter M, Payre F, Plaza S. 2010. Zona Pellucida Domain Proteins Remodel the Apical Compartment for Localized Cell Shape Changes. Dev Cell 18: 64–76.

Futahashi R, Okamoto S, Kawasaki H, Zhong YS, Iwanaga M, Mita K, Fujiwara H. 2008. Genome-wide identification of cuticular protein genes in the silkworm, Bombyx mori. Insect Biochem Mol Biol 38: 1138–1146.

Ghiradella H. 1974. Development of ultraviolet-reflecting butterfly scales: How to make an interference filter. J Morphol 142: 395–409. http://doi.wiley.com/10.1002/jmor.1051420404.

Ghiradella H. 2010. Insect Cuticular Surface Modifications: Scales and Other Structural Formations. 1st ed. Elsevier Ltd. 10.1016/S0065-2806(10)38006-4.

Ghiradella H. 1994. Structure of butterfly scales: Patterning in an insect cuticle. Microsc Res Tech 27: 429–438. 10.1002/jemt.1070270509.

Ghiradella HT, Butler MW. 2009. Many variations on a few themes : a broader look at development of iridescent scales (and feathers). *Journal of the Royal Society*, Interface / the Royal Society 6: S243–S251.

Ghosh N, Treisman JE. 2024. Apical cell expansion maintained by Dusky-like establishes a scaffold for corneal lens morphogenesis. Sci Adv 10.

Giraldo MA, Stavenga DG. 2016. Brilliant iridescence of Morpho butterfly wing scales is due to both a thin film lower lamina and a multilayered upper lamina. J Comp Physiol A Neuroethol Sens Neural Behav Physiol 202: 381–388.

Gorb SN. 2009. Functional surfaces in biology. Springer Netherlands.

Hines HM, Papa R, Ruiz M, Papanicolaou A, Wang C, Nijhout HF, McMillan WO, Reed RD. 2012. Transcriptome analysis reveals novel patterning and pigmentation genes underlying Heliconius butterfly wing pattern variation. BMC Genomics 13. 10.1186/1471-2164-13-288.

Hu Y, Comjean A, Attrill H, Antonazzo G, Thurmond J, Chen W, Li F, Chao T, Mohr SE, Brown NH, et al. 2023. PANGEA: a new gene set enrichment tool for Drosophila and common research organisms. Nucleic Acids Res 51: W419–W426. 10.1093/nar/gkad331.

Itakura Y, Wada H, Inagaki S, Hayashi S. 2026. Mechanical control of the insect extracellular matrix nanostructure. Sci Adv 12: eadw5022.

Li H, Handsaker B, Wysoker A, Fennell T, Ruan J, Homer N, Marth G, Abecasis G, Durbin R, Subgroup 1000 Genome Project Data Processing. 2009. The Sequence Alignment/Map format and SAMtools. Bioinformatics 25: 2078–2079. 10.1093/bioinformatics/btp352.

Liu J, Chen Z, Xiao Y, Asano T, Li S, Peng L, Chen E, Zhang J, Li W, Zhang Y, et al. 2021. Lepidopteran wing scales contain abundant cross-linked film-forming histidine-rich cuticular proteins. Commun Biol 4. 10.1038/s42003-021-01996-4.

Lloyd VJ, Burg SL, Harizanova J, Garcia E, Hill O, Enciso-Romero J, Cooper RL, Flenner S, Longo E, Greving I, et al. 2024. The actin cytoskeleton plays multiple roles in structural colour formation in butterfly wing scales. Nat Commun 15: 4073. 10.1038/s41467-024-48060-3.

Locke M. 1966. The structure and formation of the cuticulin layer in the epicuticle of an insect, Calpodes ethlius (Lepidoptera, Hesperiidae). J Morphol 118: 461–494.

Loh LS, DeMarr KA, Tsimba M, Heryanto C, Berrio A, Patel NH, Martin A, McMillan WO, Wray GA, Hanly JJ. 2025. Lepidopteran scale cells derive from sensory organ precursors through a canonical lineage. Development (Cambridge) 152.

Matsuoka Y, Monteiro A. 2018. Melanin Pathway Genes Regulate Color and Morphology of Butterfly Wing Scales. Cell Rep 24: 56–65.

McDougal AD, Kang S, Yaqoob Z, So PTC, Kolle M. 2021. In vivo visualization of butterfly scale cell morphogenesis in Vanessa cardui. Proceedings of the National Academy of Sciences 118: e2112009118. 10.1073/pnas.2112009118.

McKenna A, Hanna M, Banks E, Sivachenko A, Cibulskis K, Kernytsky A, Garimella K, Altshuler D, Gabriel S, Daly M, et al. 2010. The genome analysis toolkit: A MapReduce framework for analyzing next-generation DNA sequencing data. Genome Res 20: 1297– 1303.

Midic U, VandeVoort CA, Latham KE. 2018. Determination of single embryo sex in Macaca mulatta and Mus musculus RNA-Seq transcriptome profiles. Physiol Genomics 50: 628– 635. 10.1152/physiolgenomics.00001.2018.

Moussian B. 2010. Recent advances in understanding mechanisms of insect cuticle differentiation. Insect Biochem Mol Biol 40: 363–75. http://www.ncbi.nlm.nih.gov/pubmed/20347980 (Accessed April 30, 2014).

Moussian B, Seifarth C, Müller U, Berger J, Schwarz H. 2006. Cuticle differentiation during Drosophila embryogenesis. Arthropod Struct Dev 35: 137–152.

Nagaraj R, Adler PN. 2012. Dusky-like functions as a Rab11 effector for the deposition of cuticle during Drosophila bristle development. Development 139: 906–916.

Noh MY, Kramer KJ, Muthukrishnan S, Kanost MR, Beeman RW, Arakane Y. 2014. Two major cuticular proteins are required for assembly of horizontal laminae and vertical pore canals in rigid cuticle of Tribolium castaneum. Insect Biochem Mol Biol 53: 22–29. 10.1016/j.ibmb.2014.07.005.

Noh MY, Muthukrishnan S, Kramer KJ, Arakane Y. 2016. Cuticle formation and pigmentation in beetles. Curr Opin Insect Sci 17: 1–9. 10.1016/j.cois.2016.05.004.

Noh MY oung, Muthukrishnan S, Kramer KJ, Arakane Y. 2015. Tribolium castaneum RR-1 cuticular protein TcCPR4 is required for formation of pore canals in rigid cuticle. PLoS Genet 11: e1004963. 10.1371/journal.pgen.1004963.

Öztürk-Çolak A, Moussian B, Araújo SJ, Casanova J. 2016. A feedback mechanism converts individual cell features into a supracellular ECM structure in Drosophila trachea ed. U. Banerjee. Elife 5: e09373. 10.7554/eLife.09373.

Parnell AJ, Bradford JE, Curran E V., Washington AL, Adams G, Brien MN, Burg SL, Morochz C, Fairclough JPA, Vukusic P, et al. 2018. Wing scale ultrastructure underlying convergent and divergent iridescent colours in mimetic Heliconius butterflies. J R Soc Interface 15.

Perez-Riverol Y, Bandla C, Kundu DJ, Kamatchinathan S, Bai J, Hewapathirana S, John NS, Prakash A, Walzer M, Wang S, et al. 2025. The PRIDE database at 20 years: 2025 update. Nucleic Acids Res 53: D543–D553. 10.1093/nar/gkae1011.

Pertea M, Pertea GM, Antonescu CM, Chang T-C, Mendell JT, Salzberg SL. 2015. StringTie enables improved reconstruction of a transcriptome from RNA-seq reads. Nat Biotechnol 33: 290–295. 10.1038/nbt.3122.

Pinheiro J, Bates D, R-core. 2021. nlme: Linear and Nonlinear Mixed Effects Models. https://svn.r-project.org/R-packages/trunk/nlme/.

Plaza S, Chanut-Delalande H, Fernandes I, Wassarman PM, Payre F. 2010. From A to Z: Apical structures and zona pellucida-domain proteins. Trends Cell Biol 20: 524–532.

Poplin R, Chang P-C, Alexander D, Schwartz S, Colthurst T, Ku A, Newburger D, Dijamco J, Nguyen N, Afshar PT, et al. 2018. A universal SNP and small-indel variant caller using deep neural networks. Nat Biotechnol 36: 983–987. 10.1038/nbt.4235.

Prakash A, Dion E, Banerjee T Das, Monteiro A. 2024. The molecular basis of scale development highlighted by a single-cell atlas of Bicyclus anynana butterfly pupal forewings. Cell Rep 43.

Prakash A, Finet C, Banerjee T Das, Saranathan V, Monteiro A. 2022. Antennapedia and optix regulate metallic silver wing scale development and cell shape in Bicyclus anynana butterflies. Cell Rep 40.

Prakash A, Lloyd V, Yang Y, Jessop AL, McDougal A, Pirih P, Duclos G, Parnell A, Kolle M, Nadeau NJ, et al. 2026. Structure Formation in Butterfly Scales: Interplay of Genetic Control, Mechanical Instabilities, and Dynamic Material Properties. Adv Funct Mater 36.

Qiao L, Xiong G, Wang R, He S, Chen J, Tong X, Hu H, Li C, Gai T, Xin Y, et al. 2014. Mutation of a Cuticular Protein, BmorCPR2, Alters Larval Body Shape and Adaptability in Silkworm, Bombyx mori. Genetics 196: 1103–1115. 10.1534/genetics.113.158766.

R Core Team. 2021. R: A Language and Environment for Statistical Computing. https://www.r-project.org/.

Ray RP, Matamoro-Vidal A, Ribeiro PS, Tapon N, Houle D, Salazar-Ciudad I, Thompson BJ. 2015. Patterned Anchorage to the Apical Extracellular Matrix Defines Tissue Shape in the Developing Appendages of Drosophila. Dev Cell 34: 310–322. 10.1016/j.devcel.2015.06.019.

Robinson MD, McCarthy DJ, Smyth GK. 2010. edgeR: a Bioconductor package for differential expression analysis of digital gene expression data. Bioinformatics 26: 139–140. 10.1093/bioinformatics/btp616.

Robinson MD, Oshlack A. 2010. A scaling normalization method for differential expression analysis of RNA-seq data. Genome Biol 11: R25. 10.1186/gb-2010-11-3-r25.

Roch F, Alonso CR, Akam M. 2003. Drosophila miniature and dusky encode ZP proteins required for cytoskeletal reorganisation during wing morphogenesis. J Cell Sci 116: 1199– 1207.

Sahara K, Yoshido A, Traut W. 2012. Sex chromosome evolution in moths and butterflies. Chromosome Research 20: 83–94. 10.1007/s10577-011-9262-z.

Seah KS, Saranathan V. 2023. Hierarchical morphogenesis of swallowtail butterfly wing scale nanostructures eds. A. Amir and C. Desplan. Elife 12: RP89082. 10.7554/eLife.89082.

Stavenga DG, Leertouwer HL, Wilts BD. 2014. Coloration principles of nymphaline butterflies - thin films, melanin, ommochromes and wing scale stacking. J Exp Biol 217: 2171–80. http://www.ncbi.nlm.nih.gov/pubmed/24675561.

Su Y, Liu J, Liu W, Ma Y, Zhao Z, Zhao X, Zhang J. 2026. Zona pellucida proteins Piopio and Dumpy are essential for the apical extracellular matrix assembly and survival of Locusta migratoria. Pest Manag Sci 82: 7723–7735. 10.1002/ps.70835.

Surapaneni VA, Bold G, Speck T, Thielen M. 2020. Spatio-temporal development of cuticular ridges on leaf surfaces of Hevea brasiliensis alters insect attachment. R Soc Open Sci 7: 201319. 10.1098/rsos.201319.

Tajiri R. 2017. Cuticle itself as a central and dynamic player in shaping cuticle. Curr Opin Insect Sci 19: 30–35. https://www.sciencedirect.com/science/article/pii/S2214574516301651.

Thakur Y, Tevatiya S, Kumar G, Jeena M, Verma V, Dixit R, Pasi S, Eapen A, Kaur J. 2025. Panoramic view of diversity and function of cuticular proteins in insects and mosquitoes biology. Frontiers in Insect Science **Volume** 5-2025. https://www.frontiersin.org/journals/insect-science/articles/10.3389/finsc.2025.1602055.

Thayer RC, Patel NH. 2023. A meta-analysis of butterfly structural colors: their color range, distribution and biological production. Journal of Experimental Biology 226: jeb245940. 10.1242/jeb.245940.

Totz JF, McDougal AD, Wagner L, Kang S, So PTC, Dunkel J, Wilts BD, Kolle M. 2024. Cell membrane buckling governs early-stage ridge formation in butterfly wing scales. Cell Rep Phys Sci 5. 10.1016/j.xcrp.2024.102063.

Willis JH. 2010. Structural cuticular proteins from arthropods: annotation, nomenclature, and sequence characteristics in the genomics era. Insect Biochem Mol Biol 40: 189–204.

Wilts BD, Rudall PJ, Moyroud E, Gregory T, Ogawa Y, Vignolini S, Steiner U, Glover BJ. 2018. Ultrastructure and optics of the prism-like petal epidermal cells of Eschscholzia californica (California poppy). New Phytologist 219: 1124–1133.

Wilts BD, Vey AJM, Briscoe AD, Stavenga DG. 2017. Longwing (Heliconius) butterflies combine a restricted set of pigmentary and structural coloration mechanisms. BMC Evol Biol 17: 1–12.

Yadav S, Majumder A. 2021. Biomimicked hierarchical 2D and 3D structures from natural templates: applications in cell biology. Biomedical Materials 16: 062001. 10.1088/1748-605X/ac21a7.

Zhang Y, Tan Q, Lin M, Shen C, Jin L, Li G. 2023. Dusky-like Is Critical for Morphogenesis of the Cellular Protuberances and Formation of the Cuticle in Henosepilachna vigintioctopunctata. Biology (Basel) 12. https://www.mdpi.com/2079-7737/12/6/866.

