## Supplementary Figures and Tables for "Extracellular matrix properties and dynamics are important for butterfly scale nanostructure development"

**Supplementary Materials and Methods**

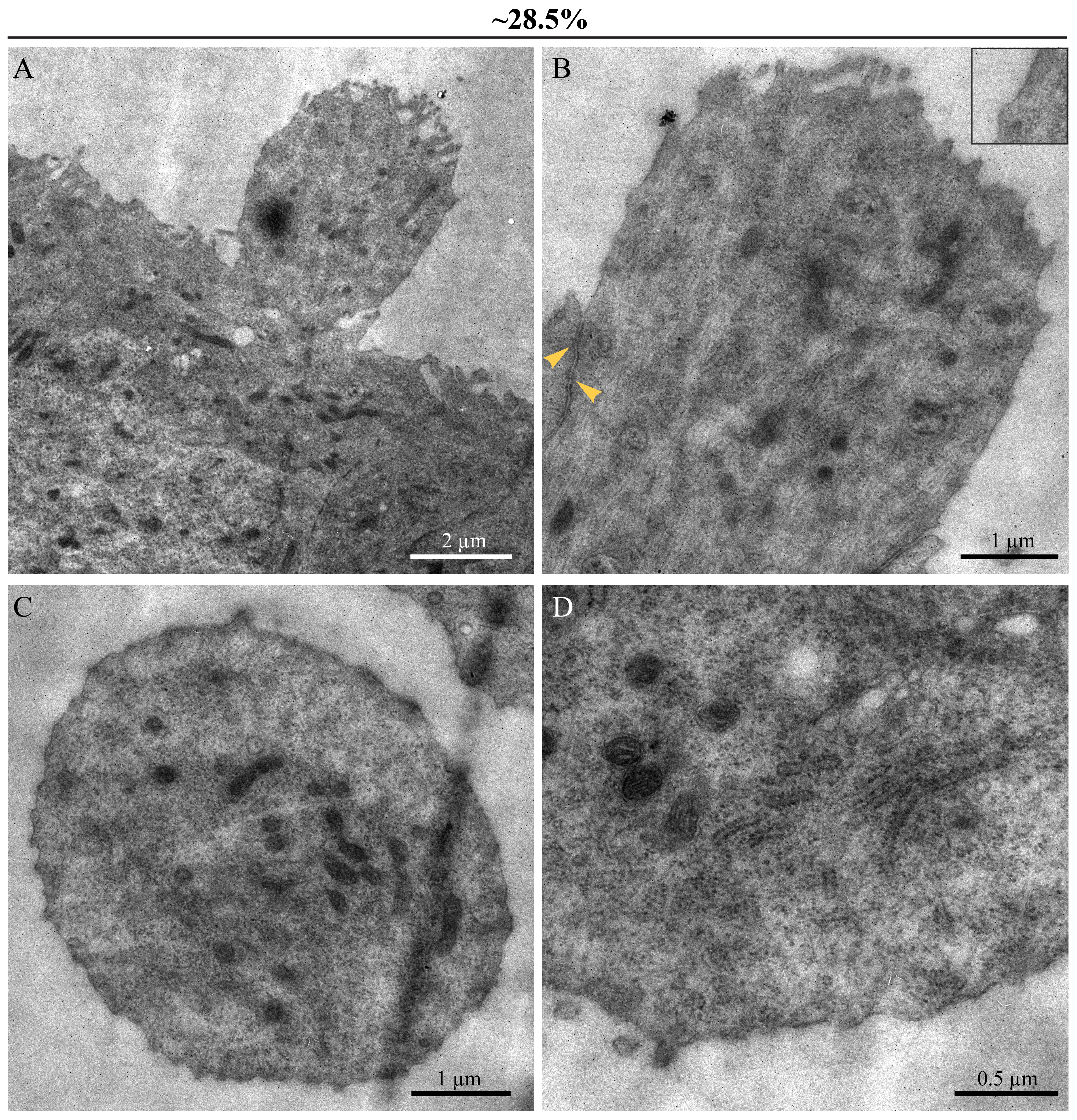

**Supplementary Figure 1: Transmission electron microscopy cross-sections of early scale buds at ~28.5 % pupal development.** (A) A longitudinal cross-section of an emerging scale. (B) Magnified view of an emerging scale. Yellow arrowheads indicate the plasma membranes of the scale and socket cell. Inset shows the scale plasma membrane and the lack of any extracellular matrix. (C-D) Transverse cross-sections of a scale.

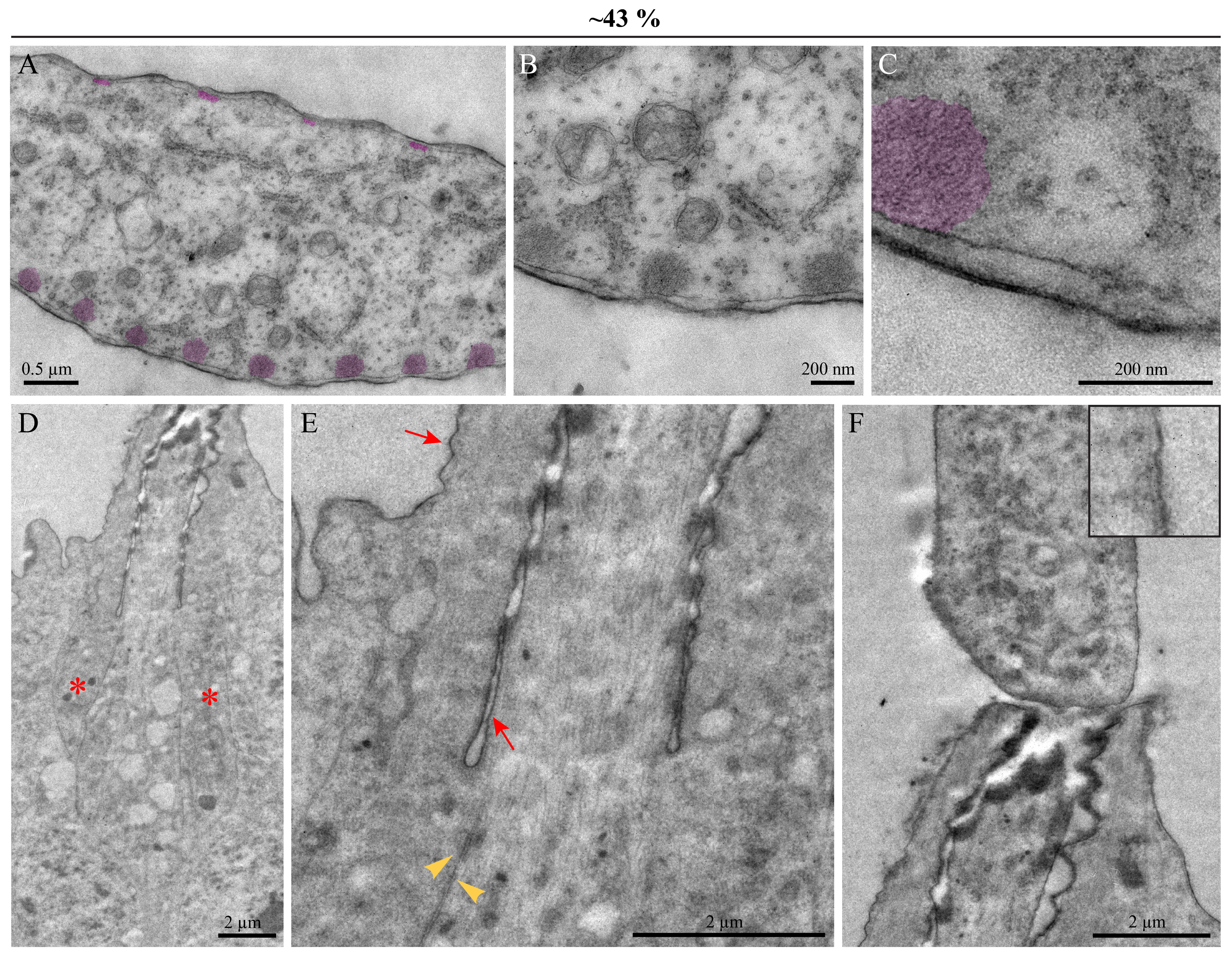

**Supplementary Figure 2: Transmission electron microscopy cross-sections of scales at ~43 % pupal development.** (A) The scale has flattened and the outermost layer of the extracellular matrix i.e., the envelope, is being secreted. The larger and smaller actin bundles on the bottom (adwing) and top (abwing) surfaces of the scale respectively are artificially colored in purple. (B, C) Magnified views of a scale transverse cross-section showing the electron dense double-layered envelope being self-assembled outside the plasma membrane. The assembly of the envelope is not uniform. Note the regions in (B) where the envelope has formed vs areas where it is still forming (fuzzy). (D) A longitudinal cross-section through the wing membrane showing the scale emerging through the socket (red stars). (E) Magnification of a region in (D) illustrating the secretion and assembly of the envelope outside the socket and scale (red arrows). Yellow arrowheads indicate the plasma membranes of the scale and socket cell. (F) A scale longitudinal cross-section. Inset shows the presence of the envelope outside the scale plasma membrane.

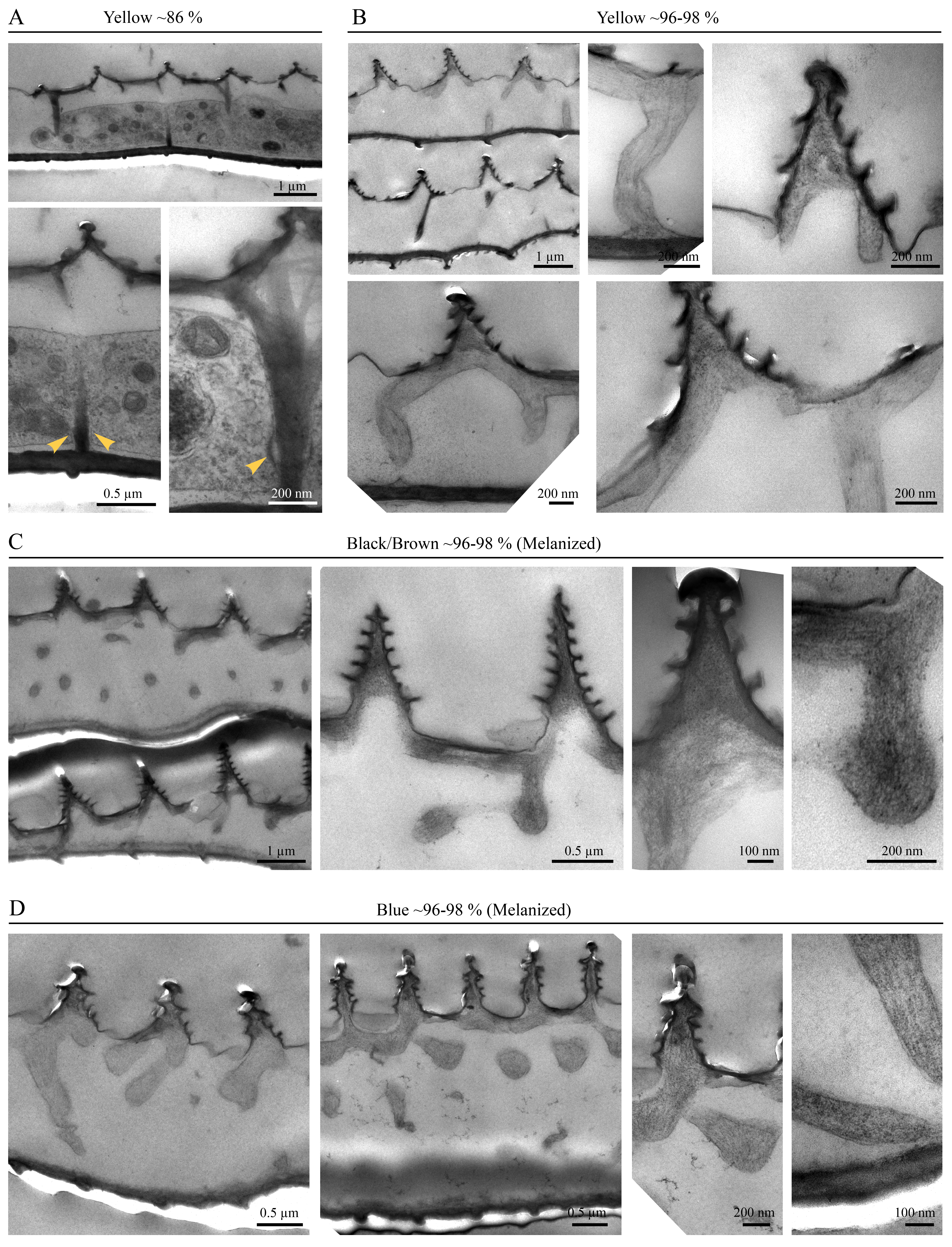

**Supplementary Figure 3: Transmission electron microscopy cross-sections of scales between ~86 - 98 % pupal development.** (A) Scales from the yellow region of the forewing at ~86% of pupal development. At this time point, the cell is retracting and trabeculae that connect the two scale surfaces are forming outside the retracting cell. Yellow arrowheads indicate the plasma membrane adjacent to the trabeculae. (B) Scales from the yellow region of the forewing at ~96-98 % of pupal development. At this stage, yellow pigments have been deposited in the wing and adjacent regions of the wing (outside the yellow patch) have become black. (C, D) Scales from the black/brown regions (dorsal/ventral respectively) (C) or blue regions (D) of the forewing at ~96-98 % of pupal development. Scales from both regions have turned dark, likely due to the deposition of melanin pigments, at this time point. Note the gradients of heavy metal stains (contrast in the images) in these scales within the ridges and in the trabeculae despite the scales being melanized and appearing visually dark.

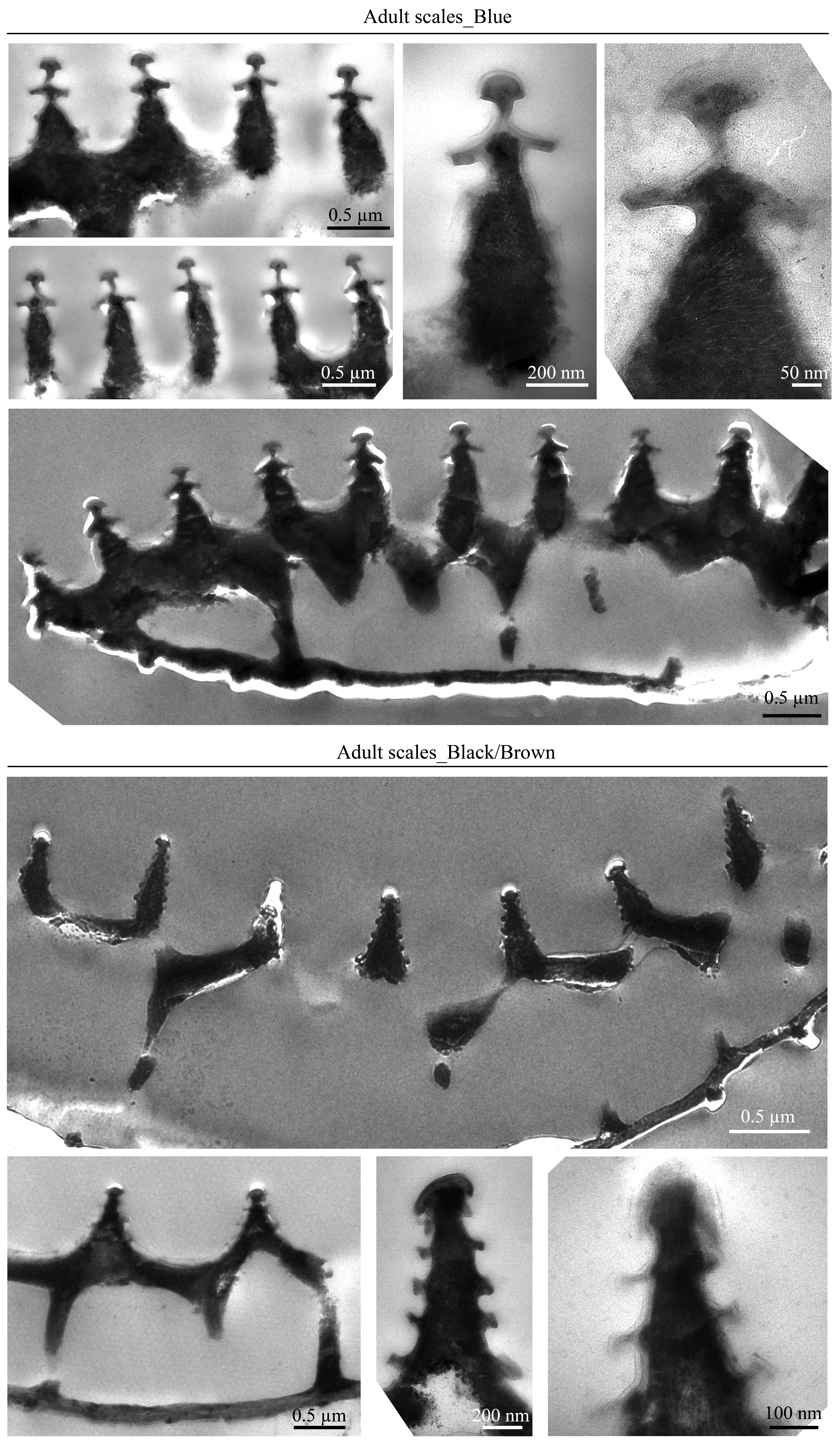
**Supplementary Figure 4: Transmission electron microscopy cross-sections of adult blue (top) and black/brown scales (bottom).**

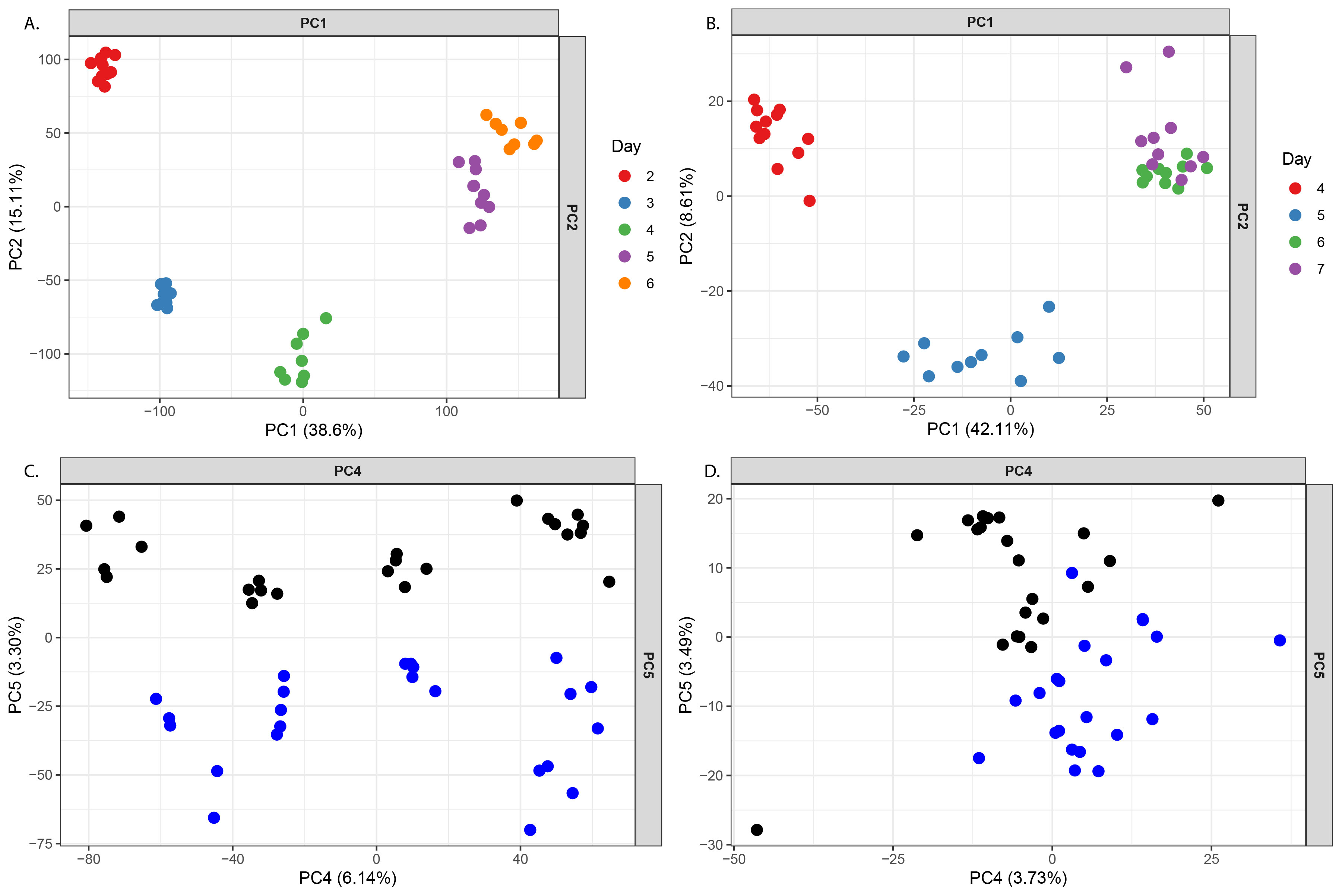

**Supplementary Figure 5: Principal component (PC) analysis of samples in the transcriptomic (A, C) and proteomic (D, B) analysis** based on normalized counts per million in the RNAseq data and normalized label-free quantification (LFQ) intensity in the proteomics data, across all genes. A and B show the first 2 PCs, with samples colored by developmental stage. C and D show the 4^th^ and 5^th^ PC with samples colored by tissue type (presumptive blue regions in blue and black regions in black).

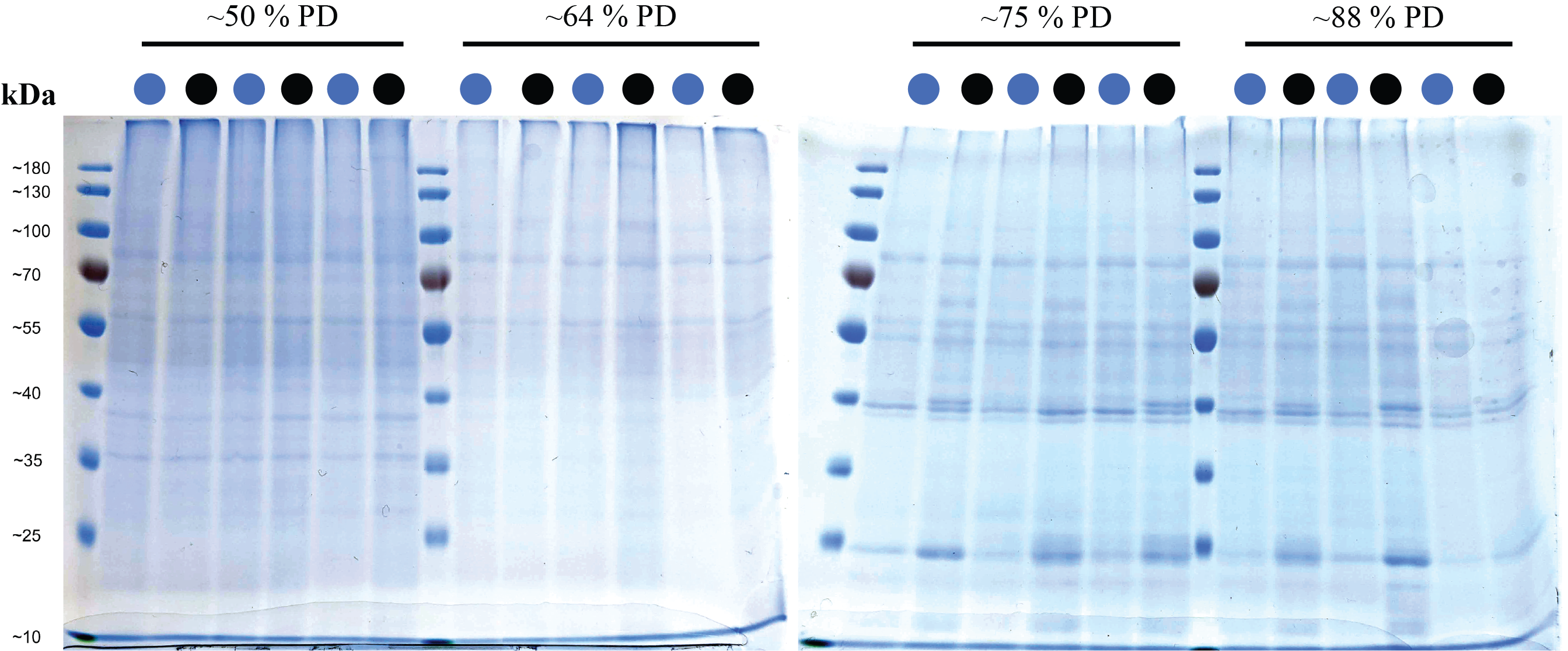

**Supplementary Figure 6: Electrophoresis protein gels at four timepoints through development.** Blue and black dots at the top indicate that the protein was extracted from presumptive blue and black wing regions. A reference size ladder is given on the left. Clearer differences in the bands present between regions can be seen at later developmental stages.

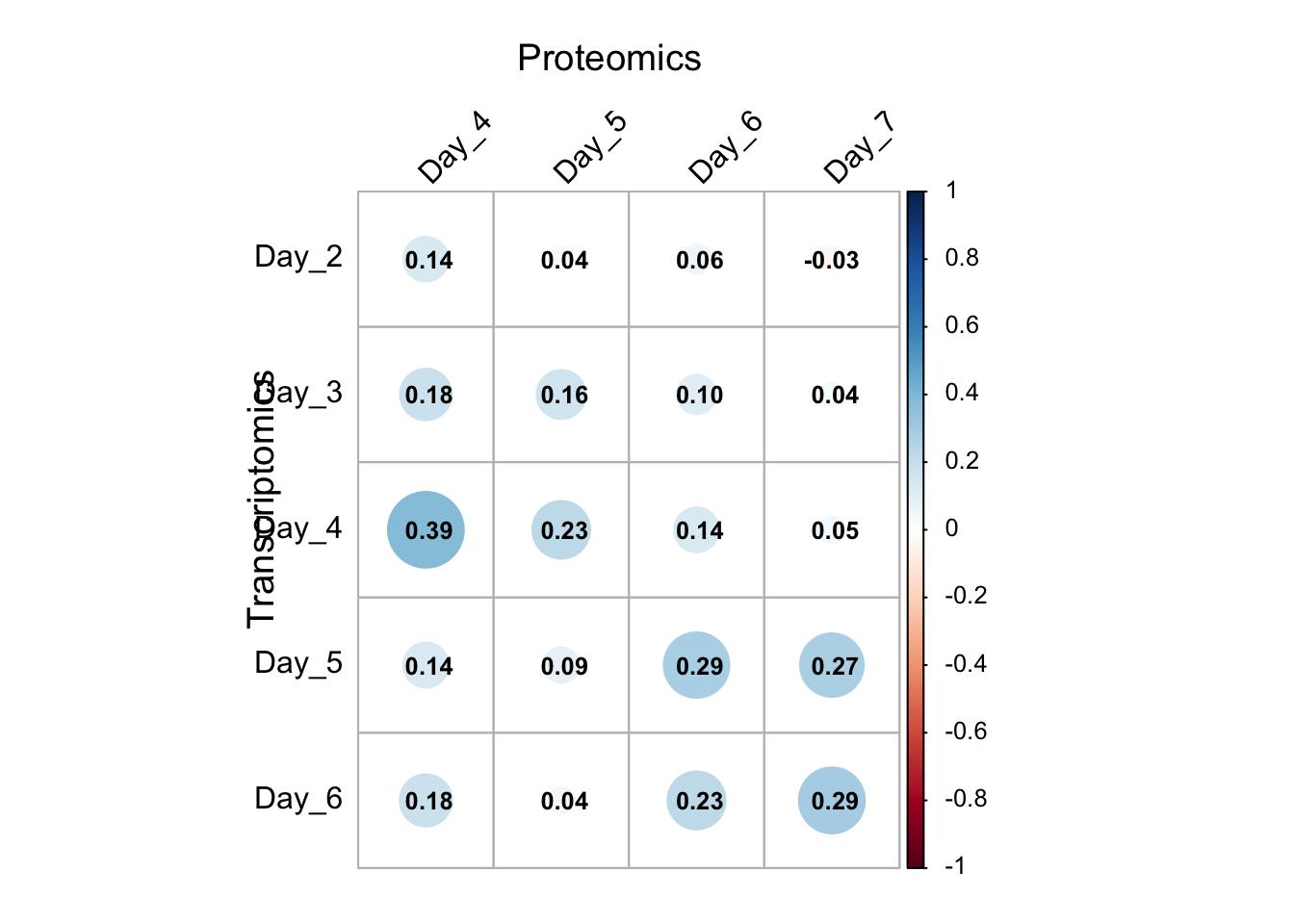

**Supplementary Figure 7: Correlations in differential expression at the gene level between the transcriptomic and proteomic data sets.** Point size and color denote the strength of the correlation. Correlation coefficients for each comparison are given. Day 2 ~ 22% PD, Day 3 ~33% PD, Day 4 = 44-50% PD, Day 5 = 56-63% PD, Day 6 = 67-75% PD, Day 7 ~ 88% PD.

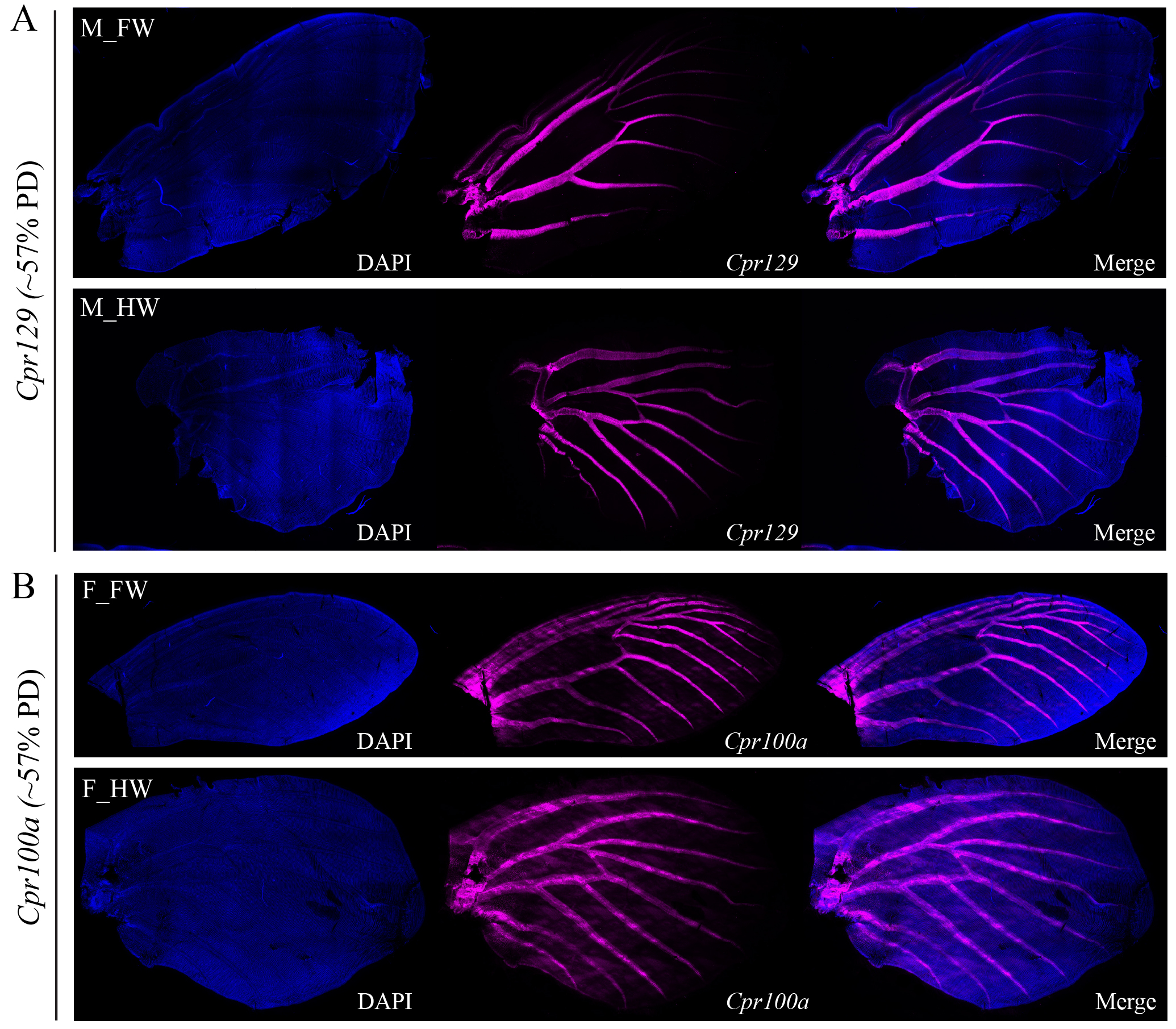

**Supplementary Figure 8: Two cuticle proteins Cpr129 and Cpr100a are expressed in the tracheal cells.** From left to right, each panel shows the nuclear staining with DAPI (blue), expression of *Cpr129* (A) or *Cpr100a* (B) mRNA (magenta) and a merge of the two on forewing (FW) and hindwing (HW) tissues at ~57 % pupal development. M = male; F = female. n = 6 wings for *Cpr129*. n = 4 wings for *Cpr100a*.

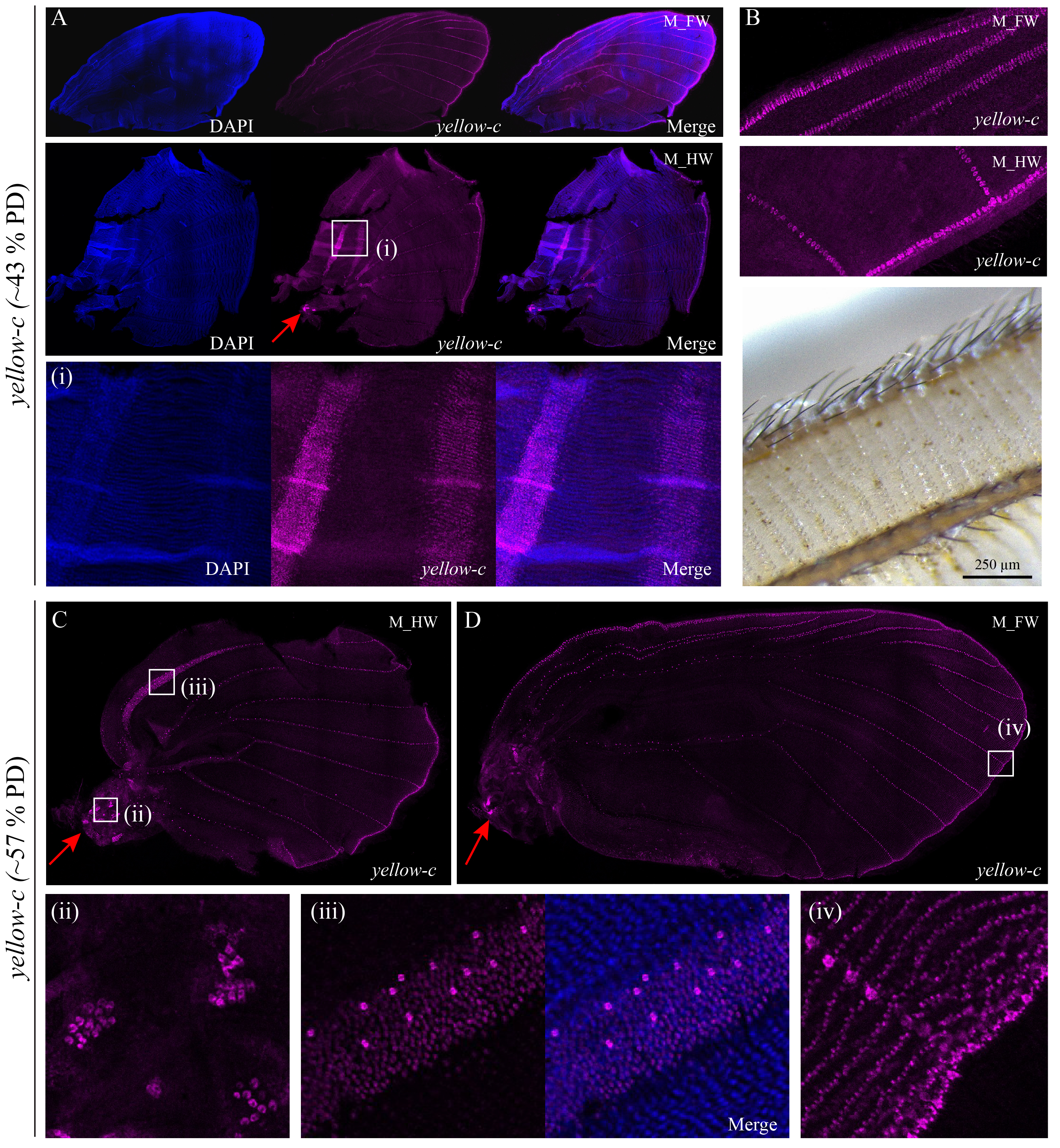

**Supplementary Figure 9: *yellow-c* is expressed in the large bristles along the veins, the androconia, and sensory organs on the wing.** (A) *yellow-c* expression on the forewing (FW) and hindwing (HW) at ~43 % pupal development (PD). The white box (i) is magnified to show expression in the androconial patch of male hindwings. The red arrow marks the expression in sensory organs at the base of the wing. (B) Magnified images of *yellow-c* expression in the bristle cells of the FW and HW. Bristles on the adult wing are shown at the bottom. (C, D) *yellow-c* expression on the HW and FW respectively at ~57 % PD. Magnified images of the (ii) sensory organs at the base of the wing (iii) expression in the androconial cells of the male HW and (iv) in the bristles along the veins and margin are shown below. Faint expression appears in scale or socket cells along the margin as seen in (iv). M = male. n = 8 wings at 43 % PD. n = 4 wings at 57 % PD.

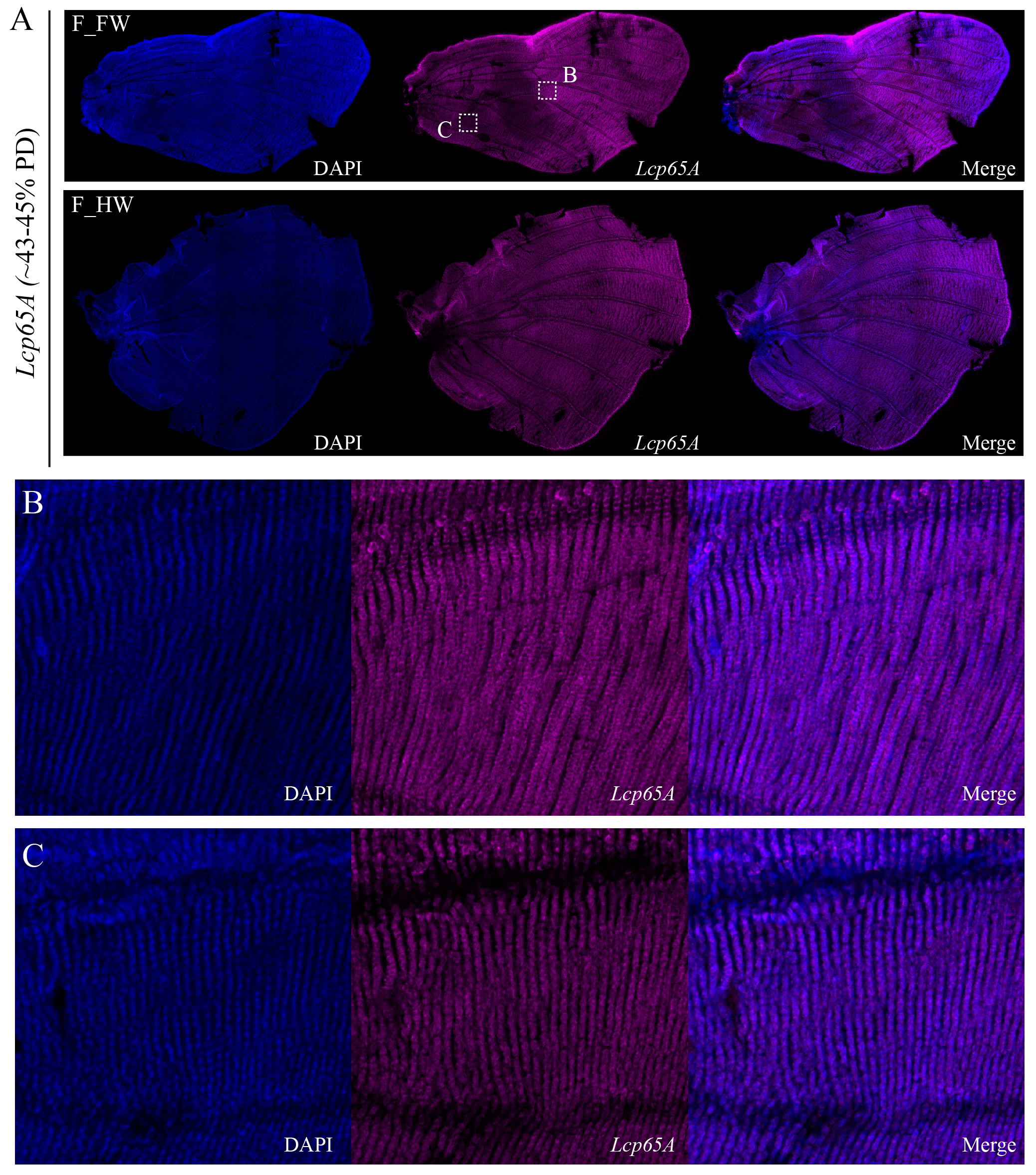

**Supplementary Figure 10: The cuticle protein *Lcp65a* has a higher mRNA expression in black scales.** (A) *Lcp65a* expression on the forewing (FW) and hindwing (HW) at ~43-45 % pupal development (PD). (B-C) Magnified images of *Lcp65a* mRNA expression in the (B) black wing region and (C) blue wing region. Panels show DAPI in blue, *Lcp65a* in magenta, and a merge of the two channels. F = female. n = 8 wings.

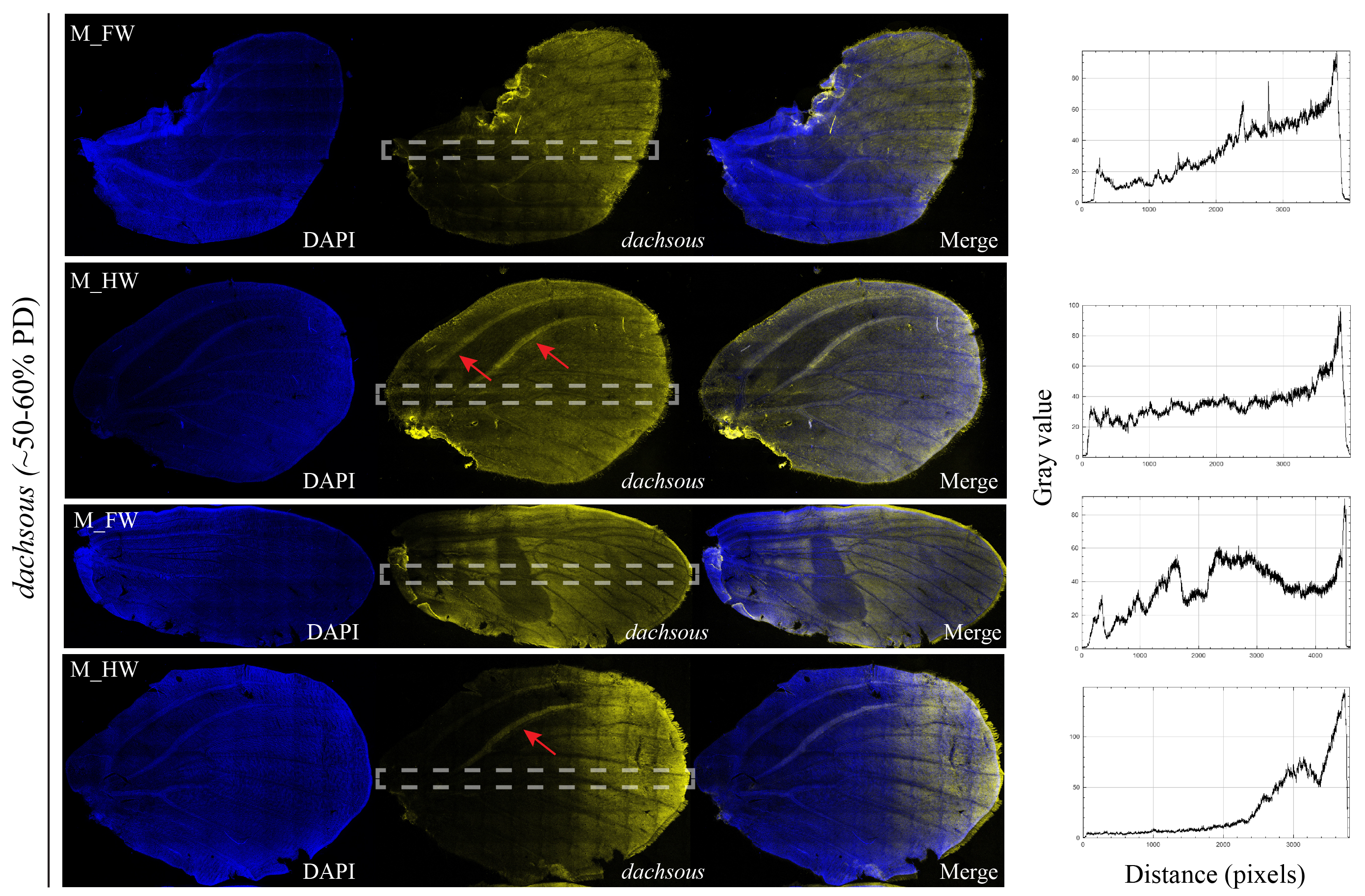

**Supplementary Figure 11: Dachsous is highly expressed in the black scales and androconia of male hindwings.** Expression of Dachsous using an anti-ds antibody in wing tissues between 50-60 % pupal development (PD). Panels on the right show pixel gray values along the region marked by the dotted white boxes. Going from left to right i.e., blue to black scales, there is an increased expression of Dachsous. Dachsous is also strongly expressed in the hindwing androconia (red arrows). n = 8 wings.

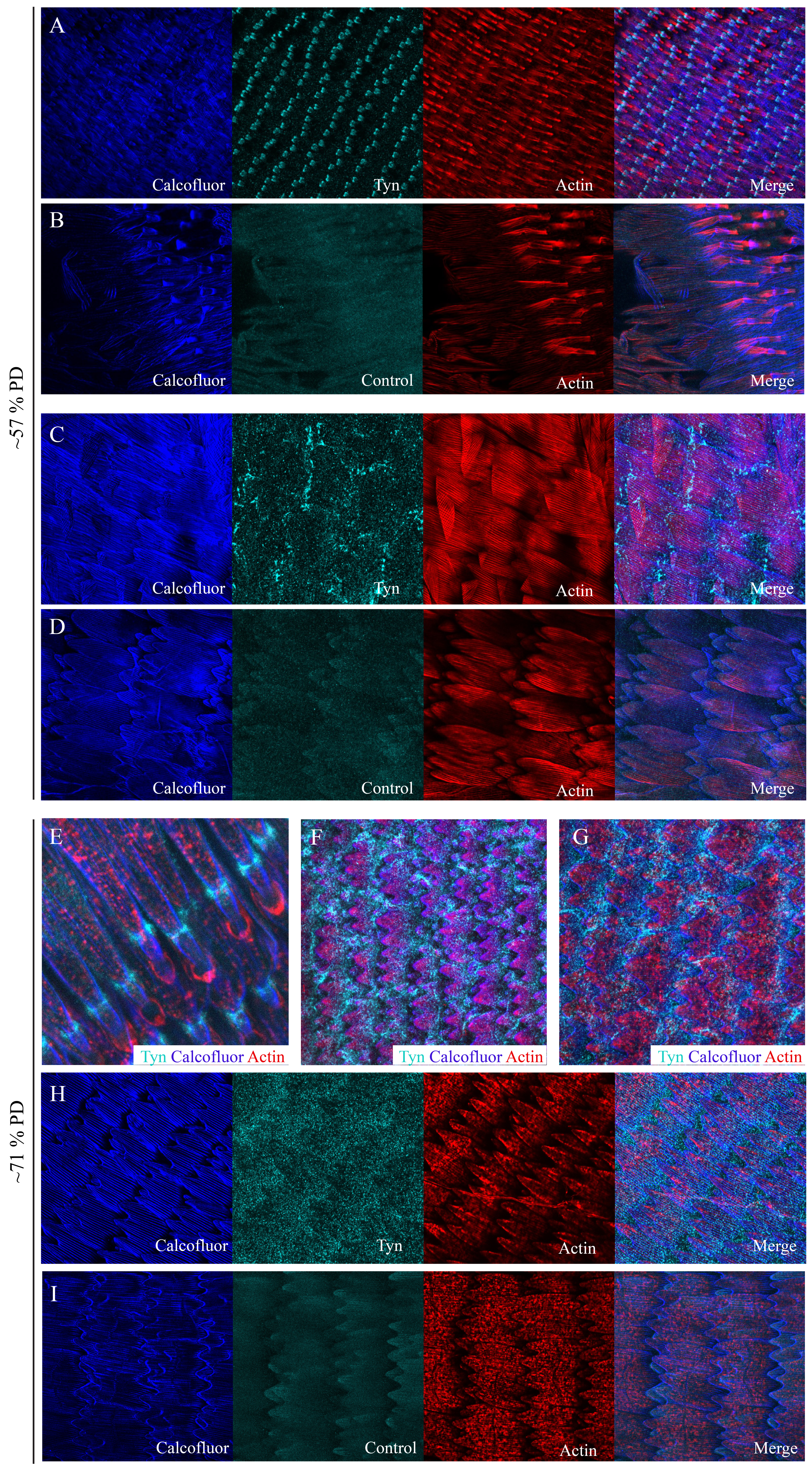

**Supplementary Figure 12: Trynity is expressed in all scale cells.** (A) At ~57 % PD Tyn is strongly expressed at the base of all scales where the scales insert into the sockets. (C) Tyn appears clustered in many small puncta over the scale blade. (B, D) Control wing images with only secondary antibodies at the respective Z planes. (E) Strong Tyn expression in a ring around the base of the scale, near the stalk, continues at ~71 % PD. (F-H) Tyn expression concentrated in many dots across the scale blade intensifies at 71 % PD. (I) Control wing with only secondary antibodies. n = 6 wings at 57 % PD. n = 4 wings at 71 % PD.

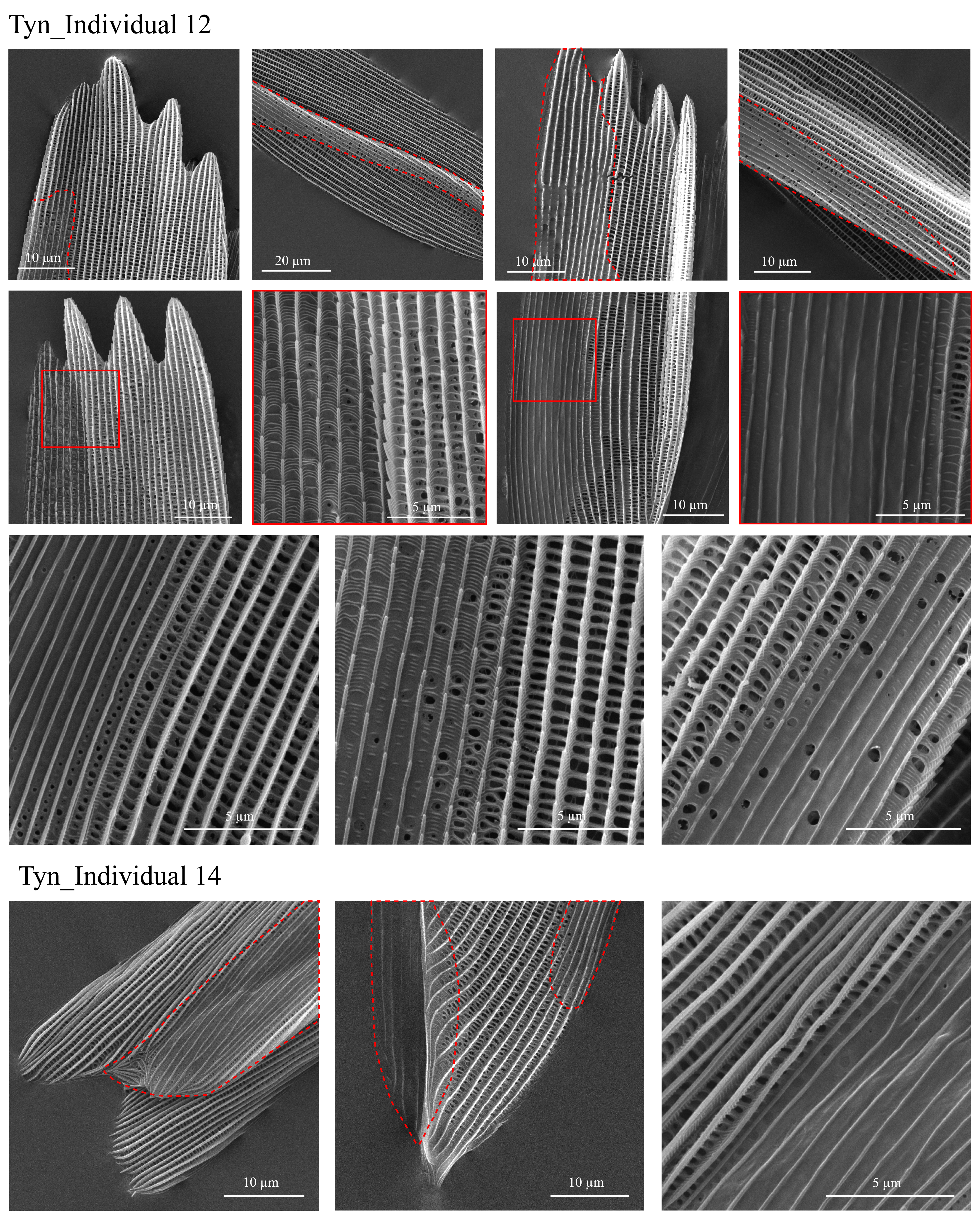

**Supplementary Figure 13: *tyn* plays a role in scale blade development.** Scanning electron micrographs of various incorrectly formed scales in two *tyn* crispants. Red dotted lines are mosaic patches where scale blades were twisted and ridges and crossribs didn’t develop as normal. Red boxes are magnified to the right.

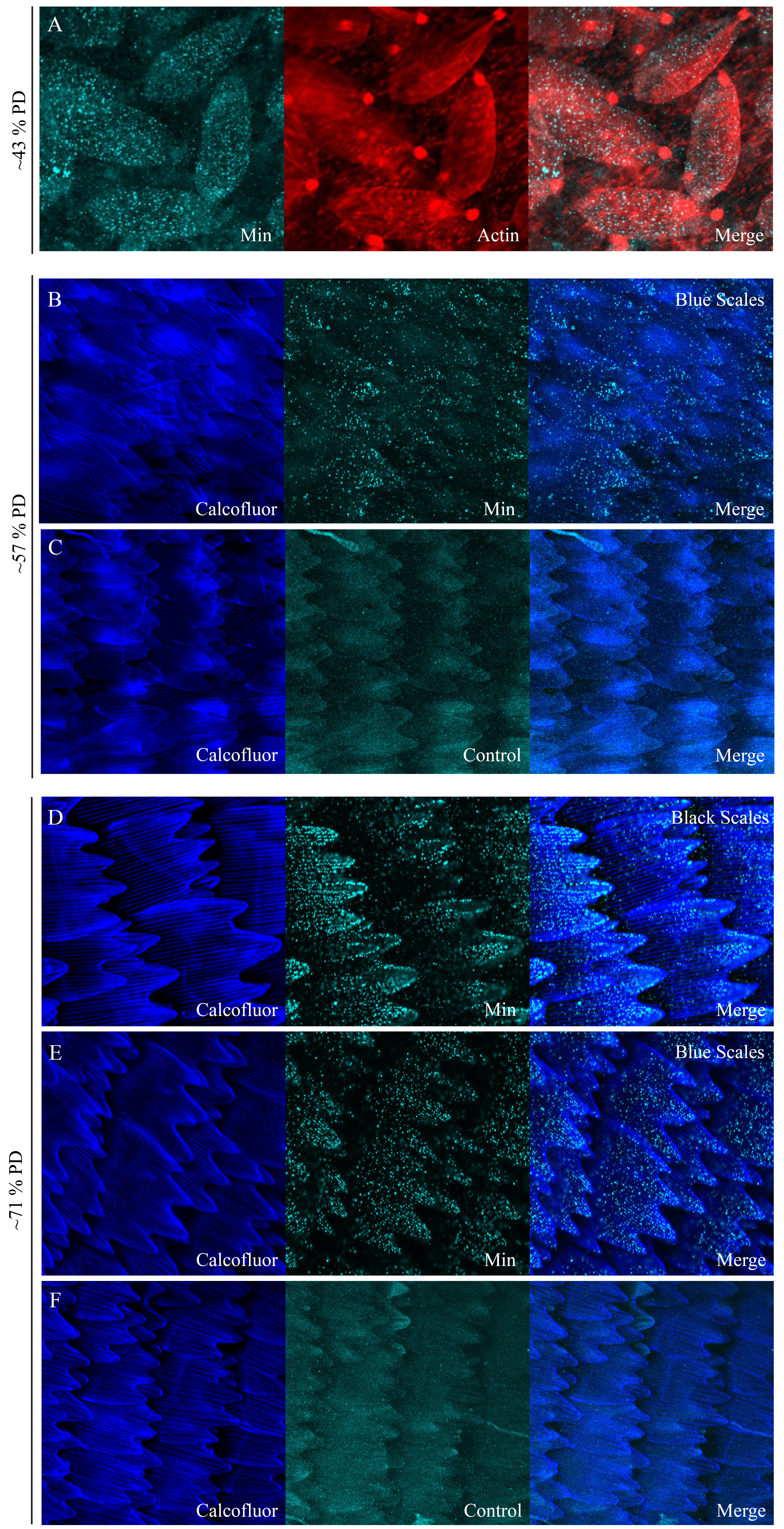

**Supplementary Figure 14: Miniature occurs in puncta in scale cells.** (A) At ~43 % PD Min occurs in small dots across the entire scale. (B-F) At the two later time points examined, Min expression becomes stronger at the distal tips of the scale, especially at the scale fingers. Expression is still seen in small puncta. (C and F) Control wing images with only secondary antibodies. n = 2 wings at 43 % PD. n = 6 wings at 57 % PD. n = 6 wings at 71 % PD.

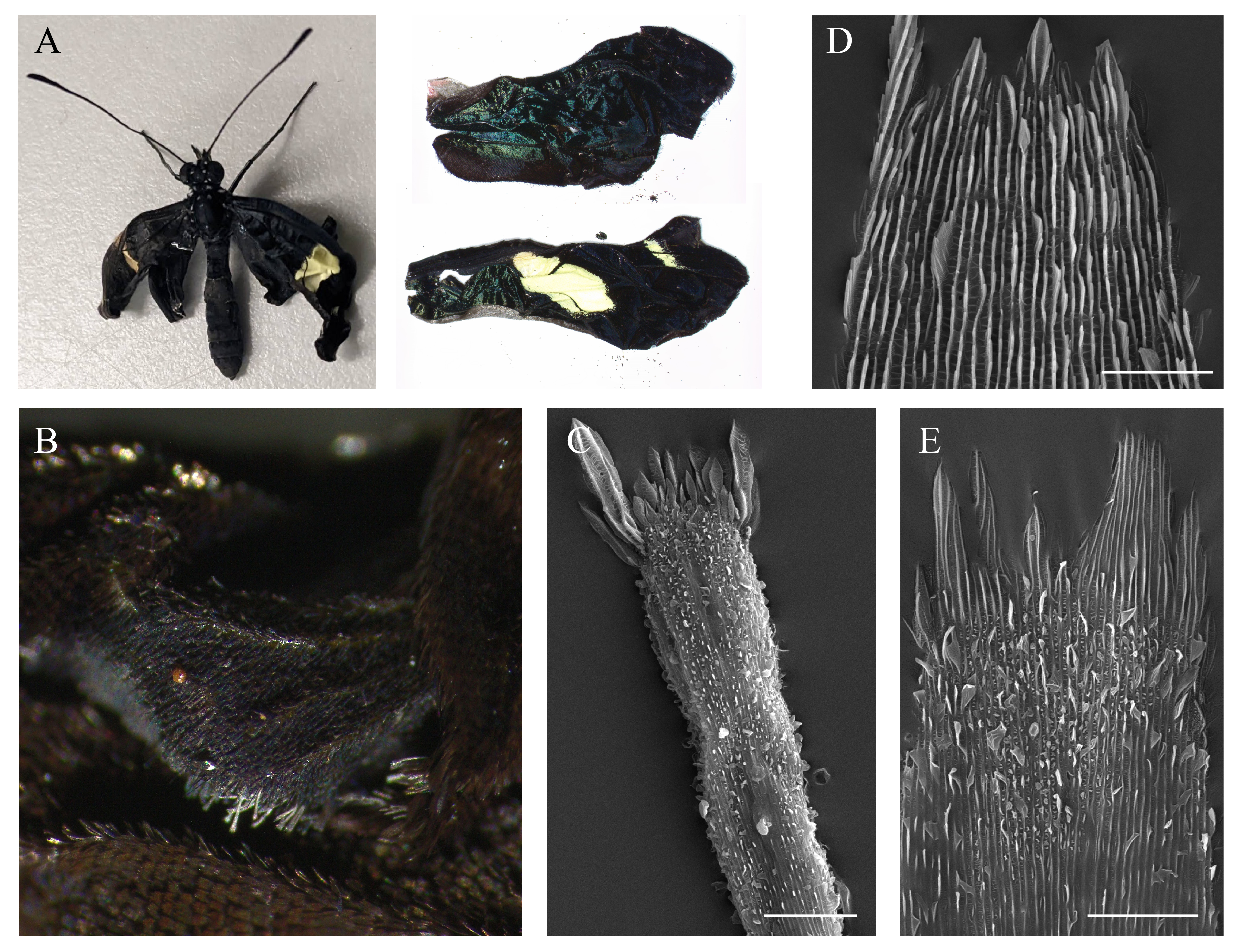

**Supplementary Figure 15: Miniature is required for wing and scale development.** (A) An eclosed *min* crispant with crumpled wings (B) Magnified view of a part of the crumpled wing with tiny and cylindrical black scales (C-E) Scanning electron micrographs of various incorrectly formed scales in *min* crispants. Scale bars are 10μm.

**Supplementary Table 1: Gene set enrichment of genes differentially expressed between presumptive blue and black wing regions at each developmental stage in the RNA-seq data.** Gene sets shown are those significant at p<0.05 after Bonferroni correction (Bonf.). The first ~9 DE genes in each category are given (followed by “…” if truncated)

| Day PD | % PD | Gene Set Category | Gene Set Name | Gene Set Size | Count Overlap Gene | % Genes in Gene Set | Fold Enrich. | P value | Bonf. | Benj. & Hoch. | Benj. & Yeku. | Overlaping Gene Symbols |
| --- | --- | --- | --- | --- | --- | --- | --- | --- | --- | --- | --- | --- |
| 2 | 0.222 | Direct GO Molecular Function | DNA-binding transcription factor activity, RNA polymerase II-specific | 252 | 18 | 12.86 | 4.52 | 0.000 | 0.000 | 0.000 | 0.000 | ara,dati,bi,Scr,tio,Poxn,hth,mirr,zfh2,dac,Antp,repo… |
| 2 | 0.222 | Direct GO Biological Processes | chitin-based cuticle development | 81 | 10 | 7.14 | 7.81 | 0.000 | 0.000 | 0.000 | 0.001 | stw,Cpr62Bc,TwdlE,Cht5,Cpr57A,Cpr65Ec,obst-B,Cpr49Aa,Cpr62Bb… |
| 2 | 0.222 | Direct GO Biological Processes | imaginal disc-derived wing morphogenesis | 108 | 11 | 7.86 | 6.45 | 0.000 | 0.001 | 0.000 | 0.001 | mirr,Bx,zfh2,Poxn,bi,hth,ap,ara,mew,hh,vg |
| 2 | 0.222 | Direct GO Cellular Component | chitin-based extracellular matrix | 57 | 8 | 5.71 | 8.88 | 0.000 | 0.002 | 0.000 | 0.003 | Cpr65Ec,TwdlE,Cpr62Bb,Cpr73D,obst-B,Cpr62Bc,Cpr49Aa,Cpr57A |
| 2 | 0.222 | Direct GO Molecular Function | structural constituent of chitin-based larval cuticle | 48 | 7 | 5.00 | 9.23 | 0.000 | 0.005 | 0.001 | 0.008 | TwdlE,Cpr62Bc,Cpr49Aa,Cpr57A,Cpr65Ec,Cpr62Bb,Cpr73D |
| 2 | 0.222 | Direct GO Molecular Function | RNA polymerase II cis-regulatory region sequence-specific DNA binding | 207 | 13 | 9.29 | 3.97 | 0.000 | 0.013 | 0.002 | 0.015 | Poxn,ems,mirr,bi,Antp,zfh2,Dll,opa,dac,grn,Optix… |
| 2 | 0.222 | Direct GO Biological Processes | regulation of transcription by RNA polymerase II | 376 | 18 | 12.86 | 3.03 | 0.000 | 0.015 | 0.002 | 0.015 | dati,knrl,bi,tio,vg,hth,ara,zfh2,opa,dac… |
| 2 | 0.222 | Direct GO Molecular Function | structural constituent of chitin-based cuticle | 13 | 4 | 2.86 | 19.47 | 0.000 | 0.023 | 0.003 | 0.019 | TwdlE,zye,obst-B,dyl |
| 2 | 0.222 | Direct GO Biological Processes | regulation of DNA-templated transcription | 319 | 16 | 11.43 | 3.17 | 0.000 | 0.025 | 0.003 | 0.019 | bi,knrl,vg,grn,ara,zfh2,ap,Scr,Poxn,hth… |
| 3 | 0.333 | Direct GO Cellular Component | chitin-based extracellular matrix | 57 | 12 | 5.58 | 8.67 | 0.000 | 0.000 | 0.000 | 0.000 | obst-A,Cpr65Ec,Cpr49Ab,Cpr49Aa,Cpr57A,obst-B,Cpr66D,Cpr49Ah… |
| 3 | 0.333 | Direct GO Molecular Function | DNA-binding transcription factor activity, RNA polymerase II-specific | 252 | 24 | 11.16 | 3.92 | 0.000 | 0.000 | 0.000 | 0.000 | ara,gl,repo,ap,Scr,dati,Dll,hth,Optix,bi,mirr,tx,unpg… |
| 3 | 0.333 | Direct GO Biological Processes | chitin-based cuticle development | 81 | 13 | 6.05 | 6.61 | 0.000 | 0.000 | 0.000 | 0.000 | Cpr66D,obst-B,Cpr49Ah,Acp65Aa,obst-A,Cpr47Ea,Cpr76Bc,Vajk4,Cpr65Ec… |
| 3 | 0.333 | Direct GO Molecular Function | structural constituent of chitin-based larval cuticle | 48 | 9 | 4.19 | 7.73 | 0.000 | 0.002 | 0.000 | 0.003 | Cpr49Aa,Cpr47Ea,Cpr57A,Cpr66D,Cpr49Ab,Acp65Aa,Cpr76Bc,Cpr49Ah,Cpr65Ec |
| 3 | 0.333 | Direct GO Cellular Component | neuron projection membrane | 19 | 6 | 2.79 | 13.01 | 0.000 | 0.003 | 0.001 | 0.005 | DIP-lambda,dpr18,DIP-kappa,DIP-gamma,dpr9,dpr6 |
| 3 | 0.333 | Direct GO Biological Processes | regulation of DNA-templated transcription | 319 | 22 | 10.23 | 2.84 | 0.000 | 0.008 | 0.001 | 0.009 | ap,knrl,tx,Poxn,grn,cas,ara,ems,unpg… |
| 3 | 0.333 | Direct GO Molecular Function | RNA polymerase II cis-regulatory region sequence-specific DNA binding | 207 | 17 | 7.91 | 3.38 | 0.000 | 0.009 | 0.001 | 0.009 | Scr,Optix,bi,svp,Dll,ci,mirr,Antp,dac,ems,toe,opa… |
| 3 | 0.333 | Direct GO Biological Processes | regulation of transcription by RNA polymerase II | 376 | 24 | 11.16 | 2.63 | 0.000 | 0.011 | 0.001 | 0.010 | knrl,dati,unpg,Dll,hth,tio,ara,gl,dac,vg… |
| 3 | 0.333 | Direct GO Biological Processes | establishment of synaptic specificity at neuromuscular junction | 4 | 3 | 1.40 | 30.90 | 0.000 | 0.046 | 0.005 | 0.035 | DIP-kappa,DIP-lambda,DIP-gamma |
| 3 | 0.333 | Direct GO Biological Processes | imaginal disc-derived wing morphogenesis | 108 | 11 | 5.12 | 4.20 | 0.000 | 0.048 | 0.005 | 0.035 | mirr,ds,tnc,Poxn,hth,ap,zfh2,ara,bi,hh,vg |
| 4 | 0.444 | Direct GO Molecular Function | structural constituent of chitin-based larval cuticle | 48 | 26 | 3.61 | 6.66 | 0.000 | 0.000 | 0.000 | 0.000 | Cpr67Fb,Cpr30F,TwdlE,Cpr62Bb,Acp65Aa,Cpr97Ea,Cpr49Ae,Cpr66Cb,Cpr65Ax2… |
| 4 | 0.444 | Direct GO Cellular Component | chitin-based extracellular matrix | 57 | 28 | 3.88 | 6.04 | 0.000 | 0.000 | 0.000 | 0.000 | Cpr66Cb,Cpr100A,obst-E,resilin,Cpr62Bc,Cpr76Bd,Cpr30F,Cpr47Ea,Cpr65Eb… |
| 4 | 0.444 | Direct GO Biological Processes | chitin-based cuticle development | 81 | 31 | 4.30 | 4.70 | 0.000 | 0.000 | 0.000 | 0.000 | Cpr97Eb,obst-B,resilin,Cpr66Cb,Cpr76Bd,Cpr49Aa,Cpr47Ea,Cpr62Bc,Cht5… |
| 4 | 0.444 | Direct GO Cellular Component | extracellular matrix | 62 | 20 | 2.77 | 3.96 | 0.000 | 0.000 | 0.000 | 0.000 | shf,resilin,Cpr76Bd,TwdlE,Cpr62Bb,Cpr30F,Cpr50Cb,Cpr66Cb,l(2)gl… |
| 4 | 0.444 | Direct GO Biological Processes | imaginal disc-derived wing morphogenesis | 108 | 26 | 3.61 | 2.96 | 0.000 | 0.001 | 0.000 | 0.001 | Fhos,zfh2,ds,vg,aos,brk,emc,hth,Bx,ara… |
| 4 | 0.444 | Direct GO Cellular Component | extracellular space | 228 | 42 | 5.83 | 2.26 | 0.000 | 0.001 | 0.000 | 0.001 | frm,Timp,hh,Pvf3,Mmp2,fj,Notum,CG9649,Lsp2,Tg… |
| 4 | 0.444 | Direct GO Cellular Component | plasma membrane | 818 | 106 | 14.70 | 1.59 | 0.000 | 0.001 | 0.000 | 0.001 | pippin,kek5,Tsp29Fb,nAChRalpha5,CCT8,Appl,Drl-2,Ctr1A,Sry-alpha… |
| 4 | 0.444 | Direct GO Cellular Component | nucleolus | 118 | 27 | 3.75 | 2.81 | 0.000 | 0.001 | 0.000 | 0.001 | Tif-IA,Mat89Ba,hoip,NO66,CrebA,Rrp5,CG32409,CG12288,Nopp140… |
| 4 | 0.444 | Direct GO Cellular Component | extracellular region | 175 | 34 | 4.72 | 2.39 | 0.000 | 0.003 | 0.000 | 0.002 | fat-spondin,Gasp,EbpIII,svr,spz4,spz6,hh,obst-E,scaf… |
| 4 | 0.444 | Direct GO Biological Processes | rRNA processing | 32 | 12 | 1.66 | 4.61 | 0.000 | 0.008 | 0.001 | 0.006 | CG9799,CG11583,ppan,CG5033,CG1542,Mat89Ba,Nop56,ais,l(3)07882… |
| 4 | 0.444 | Direct GO Biological Processes | basement membrane organization | 11 | 7 | 0.97 | 7.82 | 0.000 | 0.012 | 0.001 | 0.009 | Col4a1,Mp,Mmp2,LanB2,Pi4KIIIalpha,Ndg,Timp |
| 4 | 0.444 | Direct GO Cellular Component | small-subunit processome | 24 | 10 | 1.39 | 5.12 | 0.000 | 0.017 | 0.001 | 0.012 | l(2)k09022,U3-55K,Mat89Ba,hoip,l(2)05287,Nop56,Sas10,l(3)07882,CG9799… |
| 4 | 0.444 | Direct GO Molecular Function | RNA polymerase II cis-regulatory region sequence-specific DNA binding | 207 | 35 | 4.85 | 2.08 | 0.000 | 0.049 | 0.004 | 0.031 | gl,Mitf,bi,Hr3,Eip75B,Antp,ci,ham,slbo,ems,SoxN… |
| 5 | 0.556 | Direct GO Cellular Component | chitin-based extracellular matrix | 57 | 29 | 5.70 | 8.86 | 0.000 | 0.000 | 0.000 | 0.000 | Cpr49Ae,Cpr65Av,Lcp65Ac,Gasp,resilin,Cpr65Ec,Cpr92A,obst-B,Cpr65Ax2… |
| 5 | 0.556 | Direct GO Molecular Function | structural constituent of chitin-based larval cuticle | 48 | 26 | 5.11 | 9.43 | 0.000 | 0.000 | 0.000 | 0.000 | Cpr64Ab,Cpr67Fb,Cpr49Aa,Cpr49Ah,Acp65Aa,Ccp84Ad,Cpr62Bb,Cpr66Cb,Cpr100A… |
| 5 | 0.556 | Direct GO Biological Processes | chitin-based cuticle development | 81 | 32 | 6.29 | 6.88 | 0.000 | 0.000 | 0.000 | 0.000 | Cpr50Cb,Acp65Aa,Edg84A,Cpr65Av,obst-A,Cpr62Bb,Cpr66Cb,Cpr97Ea,Cpr49Ae… |
| 5 | 0.556 | Direct GO Cellular Component | extracellular matrix | 62 | 23 | 4.52 | 6.46 | 0.000 | 0.000 | 0.000 | 0.000 | Cpr67B,Cpr30F,resilin,Cpr62Bb,Edg84A,l(2)gl,Mp,tnc,Mmp2… |
| 5 | 0.556 | Direct GO Cellular Component | extracellular space | 228 | 38 | 7.47 | 2.90 | 0.000 | 0.000 | 0.000 | 0.000 | modSP,Amy-p,scaf,sona,Nep2,tnc,NLaz,Sod3,Phk-3,Tk,Lsp2… |
| 5 | 0.556 | Direct GO Cellular Component | extracellular region | 175 | 30 | 5.89 | 2.98 | 0.000 | 0.000 | 0.000 | 0.000 | Npc2g,PGRP-SA,scaf,gbb,GNBP1,EbpIII,Wnt6,Sema2a,hh,ChLD3… |
| 6 | 0.667 | Direct GO Molecular Function | structural constituent of chitin-based larval cuticle | 48 | 28 | 3.72 | 6.87 | 0.000 | 0.000 | 0.000 | 0.000 | Cpr49Aa,Cpr49Ac,Lcp65Ad,Cpr73D,Cpr50Cb,Cpr65Eb,Ccp84Ad,Cpr100A,TwdlE… |
| 6 | 0.667 | Direct GO Cellular Component | chitin-based extracellular matrix | 57 | 30 | 3.99 | 6.20 | 0.000 | 0.000 | 0.000 | 0.000 | resilin,Cpr65Ec,Ccp84Ad,Cpr64Ab,Cpr67B,Cpr100A,Cpr47Ea,Acp65Aa,Edg84A… |
| 6 | 0.667 | Direct GO Cellular Component | extracellular matrix | 62 | 31 | 4.12 | 5.89 | 0.000 | 0.000 | 0.000 | 0.000 | fat-spondin,frac,Ccp84Ad,shf,Tsp,Mmp1,Cpr67B,SPARC,Cpr50Cb… |
| 6 | 0.667 | Direct GO Biological Processes | chitin-based cuticle development | 81 | 35 | 4.65 | 5.09 | 0.000 | 0.000 | 0.000 | 0.000 | Cpr49Ac,Cpr62Bc,Cpr66D,Cpr97Eb,Cpr50Cb,Lcp65Ac,Cpr73D,Cpr65Ax2,stw,Cht2… |
| 6 | 0.667 | Direct GO Cellular Component | extracellular space | 228 | 54 | 7.18 | 2.79 | 0.000 | 0.000 | 0.000 | 0.000 | Argk1,Ace,Actbeta,Tg,GLaz,sona,Adgf-A,Eh,Wnt5,Spn27A… |
| 6 | 0.667 | Direct GO Cellular Component | apical plasma membrane | 73 | 23 | 3.06 | 3.71 | 0.000 | 0.000 | 0.000 | 0.000 | Ca-alpha1D,rols,tyn,rst,Np,osy,dome,promL,sns,scaf… |
| 6 | 0.667 | Direct GO Biological Processes | muscle contraction | 8 | 7 | 0.93 | 10.31 | 0.000 | 0.000 | 0.000 | 0.001 | Mlc1,up,Mhc,RyR,wupA,Tm1,Ca-alpha1D |
| 6 | 0.667 | Direct GO Biological Processes | basement membrane organization | 11 | 8 | 1.06 | 8.57 | 0.000 | 0.001 | 0.000 | 0.001 | Pi4KIIIalpha,Mmp2,Mp,vkg,Ndg,LanB2,Col4a1,Mmp1 |
| 6 | 0.667 | Direct GO Cellular Component | extracellular region | 175 | 35 | 4.65 | 2.36 | 0.000 | 0.003 | 0.000 | 0.002 | Wnt5,Cpr49Ac,Phk-3,tnc,Wnt10,resilin,CG13675,yellow-h,dpr6 … |
| 6 | 0.667 | Direct GO Biological Processes | imaginal disc-derived wing morphogenesis | 108 | 25 | 3.32 | 2.73 | 0.000 | 0.006 | 0.001 | 0.005 | emc,hth,ast,ara,mys,crol,mam,aos,mthl1… |
| 6 | 0.667 | Direct GO Cellular Component | plasma membrane | 818 | 105 | 13.96 | 1.51 | 0.000 | 0.013 | 0.001 | 0.010 | osy,Dscam1,NtR,mthl4,Eip63F-1,kst,OtopLc,Reck,Cad89D… |
| 6 | 0.667 | Direct GO Molecular Function | calcium ion binding | 133 | 27 | 3.59 | 2.39 | 0.000 | 0.032 | 0.003 | 0.022 | Cad89D,Cadps,rho-4,Mctp,scf,SPARC,Duox,rdgC,Mlc2,Dgk,tok… |
| 6 | 0.667 | Direct GO Biological Processes | cell adhesion mediated by integrin | 16 | 8 | 1.06 | 5.89 | 0.000 | 0.039 | 0.003 | 0.023 | Abi,SCAR,LanB2,mys,scb,Hem,Tsp,ItgaPS4 |
| 6 | 0.667 | Direct GO Biological Processes | cortical actin cytoskeleton organization | 25 | 10 | 1.33 | 4.71 | 0.000 | 0.040 | 0.003 | 0.023 | Prm,Abi,mwh,Hem,loco,moody,Rac1,Sbf,shi,SCAR |

**Supplementary Table 2: Gene set enrichment of proteins differentially expressed between presumptive blue and black wing regions at each developmental stage in the proteomics data.** Gene sets shown are those significant at p<0.05 after Bonferroni correction (Bonf.). The first 9 DE genes in each category are given

| Day PD | % PD | Gene Set Category | Gene Set Name | Gene Set Size | Count Overlap Gene | % Genes in Gene Set | Fold Enrich. | P value | Bonf. | Benj. & Hoch. | Benj. & Yeku. | Overlaping Gene Symbols |
| --- | --- | --- | --- | --- | --- | --- | --- | --- | --- | --- | --- | --- |
| 4 | 0.5 | Direct GO Cellular Component | basement membrane | 5 | 3 | 6.98 | 30.01 | 0.000 | 0.014 | 0.014 | 0.085 | LanB2,trol,vkg |
| 4 | 0.5 | Direct GO Biological Processes | innate immune response | 7 | 3 | 6.98 | 21.44 | 0.000 | 0.049 | 0.025 | 0.144 | Tep3,Tg,spz6 |
| 5 | 0.625 | Direct GO Biological Processes | defense response to Gram-positive bacterium | 2 | 2 | 6.25 | 67.22 | 0.000 | 0.034 | 0.034 | 0.193 | Tep3,PPO2 |
| 6 | 0.75 | Direct GO Biological Processes | chitin-based cuticle development | 42 | 20 | 6.10 | 3.12 | 0.000 | 0.001 | 0.001 | 0.006 | Cpr49Ah,Cpr66Cb,Cpr62Bc,Ccp84Ag,mmy,Dhc64C,Cpr100A,Idgf6… |
| 6 | 0.75 | Direct GO Molecular Function | structural constituent of chitin-based larval cuticle | 21 | 13 | 3.96 | 4.06 | 0.000 | 0.002 | 0.001 | 0.007 | Cpr97Ea,Cpr62Bc,Lcp65Ac,Cpr49Ah,Ccp84Ag,Cpr100A,Cpr64Ab… |
| 6 | 0.75 | Direct GO Cellular Component | extracellular matrix | 26 | 14 | 4.27 | 3.53 | 0.000 | 0.007 | 0.002 | 0.018 | dpy,Cpr64Ac,Hml,l(2)gl,Ccp84Ad,Cpr62Bc,Cpr66Cb,Cpr76Bb,CG15080… |
| 6 | 0.75 | Direct GO Cellular Component | chitin-based extracellular matrix | 28 | 14 | 4.27 | 3.28 | 0.000 | 0.021 | 0.005 | 0.041 | Cpr76Bb,Cpr62Bc,Ccp84Ad,Lcp65Ac,Cpr49Aa,Cpr49Ah,Gasp,Cpr64Ab,Cpr97Ea… |
| 7 | 0.875 | Direct GO Cellular Component | extracellular matrix | 26 | 8 | 7.92 | 6.55 | 0.000 | 0.008 | 0.008 | 0.059 | Plod,CG15080,Cpr64Ab,Cpr76Bb,Cpr62Bb,Cpr66Cb,l(2)gl,Cpr62Bc |
| 7 | 0.875 | Direct GO Molecular Function | structural constituent of chitin-based larval cuticle | 21 | 7 | 6.93 | 7.10 | 0.000 | 0.017 | 0.008 | 0.059 | Cpr66Cb,Cpr49Ah,Cpr64Ab,Cpr76Bb,Cpr62Bb,Lcp65Ac,Cpr62Bc |
| 7 | 0.875 | Direct GO Biological Processes | dorsal closure | 30 | 8 | 7.92 | 5.68 | 0.000 | 0.027 | 0.009 | 0.063 | cora,Inx3,ed,Ack,mys,shot,Nrx-IV,l(2)gl |

**Supplementary Table 3: Primers for HCR – Excel file**

**Supplementary Table 4: Hybridization chain reaction (HCR3.0 – Fluorescent based in situ hybridization) buffers**

| **Buffer** | **Composition** | **Amount** |
| --- | --- | --- |
| 1X PBST (50 mL)  * Sterilize by autoclaving | 1X PBS | 5 mL of 10X PBS |
|  | 0.1% Tween 20 | 50 μl of Tween 20 |
|  | DEPC treated water | To 50 mL |
| 20X SSC (500 mL)  *Adjust the pH to 7.0 with 1M HCl and sterilize by autoclaving. | NaCl | 87.7g |
|  | Sodium citrate | 44.1g |
|  | DEPC treated water | To 500 mL |
| 5X SSCT (40 mL) | 5X SSC | 10 mL of 20X SSC |
|  | 0.1% Tween 20 | 40 μl of Tween 20 |
|  | DEPC treated water | To 40 mL |
| Hybridization buffer (40 mL)  *Store at -20°C | 30% Formamide | 12 mL Formamide |
|  | 5X SSC | 10 mL of 20X SSC |
|  | 9 mM citric acid (pH 6.0) | 360 μl of 1M citric acid |
|  | 0.1% Tween 20 | 40 μl of Tween 20 |
|  | 50 μg/mL heparin | 200 μl of 10 mg/mL heparin |
|  | 1X Denhardt’s solution | 800 μl of 50X Denhardt’s solution |
|  | 5% dextran sulfate | 4 mL of 50% dextran sulfate |
|  | DEPC treated water | To 40 mL |
| Wash buffer (40 mL)  *Store at -20°C | 30% Formamide | 12 mL Formamide |
|  | 5X SSC | 10 mL of 20X SSC |
|  | 9 mM citric acid (pH 6.0) | 360 μl of 1M citric acid |
|  | 0.1% Tween 20 | 40 μl of Tween 20 |
|  | 50 μg/mL heparin | 200 μl of 10 mg/mL heparin |
|  | DEPC treated water | To 40 mL |
| Amplification buffer (40 mL) *Store at 4°C | 5X SSC | 10 mL of 20X SSC |
|  | 0.1% Tween 20 | 40 μl of Tween 20 |
|  | 5% dextran sulfate | 4 mL of 50% dextran sulfate |
|  | DEPC treated water | To 40 mL |
| Detergent solution (50 mL) *Store at 4°C | 1% SDS | 5 mL of 10% SDS (filtered) |
|  | 0.5% Tween 20 | 250 μl of Tween 20 |
|  | 50 mM Tris-HCl (pH 7.5) | 2.5 mL of 1M Tris-HCl |
|  | 1 mM EDTA (pH 8.0) | 100 μl of 0.5M EDTA |
|  | 150 mM NaCl | 1.5 mL of 5M NaCl |
|  | DEPC treated water | To 50 mL |

**Supplementary Table 5: Guide RNA sequences and genotyping primers**

| **Gene** | **Primer Number** | **Primer Sequence** |
| --- | --- | --- |
| *miniature* CRISPR Guide 1 | NN 13 | 5’GAAATTAATACGACTCACTATAGGCGGTACGTACACGCATCGGTTTTAGAGCTAGAAATAGC 3’ |
| *miniature* genotyping primers for Guide 1 | NN 18  NN 15 | Forward 5’ CAGTAACAAACCTCACATATATCG 3’  Reverse 5’ CTGCAATCAAAACCTACTTAGAG 3’ |
| *miniature* CRISPR Guide 2 | NN 20 | 5’GAAATTAATACGACTCACTATAGGGACGAGGCGAAGAGGCTGGTTTTAGAGCTAGAAATAGC 3’ |
| *miniature* genotyping primers for Guide 2 | NN 21  NN 22 | Forward 5’ GGCTGGCCTCAATTTATAC 3’  Reverse 5’ CTATTCCAGATCTACACGG 3’ |
| *miniature* CRISPR Guide 3 | NN 23 | 5’GAAATTAATACGACTCACTATAGGTGTCGGGCATCGTGCCTTTGTTTTAGAGCTAGAAATAGC 3’ |
| *miniature* genotyping primers for Guide 3 | NN 24  NN 25 | Forward 5’ CGGAATCGAACCCACAATC 3’  Reverse 5’ CTTCATATGAACAGGGGAC 3’ |
| *trinity* CRISPR Guide | NN 35 | 5’GAAATTAATACGACTCACTATAGGCCACCACACCCTACCCGAGTTTTAGAGCTAGAAATAGC 3’ |
| *trynity* genotyping primers | NN 36  NN 37 | Forward 5’ GCCTTGTTTCTTGCCAGG 3’  Reverse 5’ AAATTGTTACTTACTTTCGCAG 3’ |
| *CPR128* CRISPR Guide | NN 32 | 5’GAAATTAATACGACTCACTATAGGTGTCAATGATTTTCACACGTTTTAGAGCTAGAAATAGC 3’ |
| *CPR128* genotyping primers | NN 47  NN 48 | Forward 5’ TATATAATGGGACGCGCGGG 3’  Reverse 5’ GCTTTGTTTTGCTATGGAAGGC 3’ |

**Supplementary Table 6:** **CRISPR/Cas9 injection concentrations and mutation frequencies**

| Guide | Cas9 Protein Conc (ng/µl) | Guide RNA Conc (ng/µl) | Eggs injected | Eggs hatched | Hatch ratio | Total adults | Sequenced | Crispants |
| --- | --- | --- | --- | --- | --- | --- | --- | --- |
| *miniature* Guide 1 | 500 | 300 | 283 | 11 | 3.88 % | 8 | 4 | 3 |
| *miniature* Guide 1 | 300 | 150 | 270 | 25 | 9.25 % | 3 Adults  2 Pupae with malformed wings | 2 | 2 |
| *miniature* Guide 2 | 400 | 200 | 443 | 36 | 8.13 % | 5 | 5 | 2 |
| *miniature* Guide 3 | 300 | 150 | 496 | 34 | 6.85 % | 13 | 13 | 2 |
| *CPR128* | 500 | 300 | 159 | 50 | 31.44 % | 40 | 30 | 17 |
| *CPR128* | 300 | 150 | 115 | 27 | 23.47 % |  |  |  |
| *CPR128* | 500 | 300 | 145 | 9 | 6.2 % |  |  |  |
| Control (washed in Milton’s solution but not injected) | - | - | 42 | 13 | 30.9 % | 10 | - | - |
